# Methylmercury demethylation and volatilization by animals expressing microbial enzymes

**DOI:** 10.1101/2023.12.11.571038

**Authors:** K Tepper, J King, PM Cholan, C Pfitzner, M Morsch, SC Apte, M Maselko

**Affiliations:** Applied Biosciences, Macquarie University; Sydney, NSW 2109, Australia; CSIRO Environment, Lucas Heights, Sydney; NSW 2234, Australia; Faculty of Medicine, Health and Human Sciences, Macquarie Medical School, Macquarie University; Sydney, NSW 2109, Australia; Biomolecular Discovery Research Centre, Macquarie University; Sydney, NSW 2109, Australia; ARC Centre of Excellence in Synthetic Biology, Macquarie University; Sydney, NSW 2109, Australia

## Abstract

Methylmercury is a highly toxic pollutant that accumulates in food webs where it is inaccessible to current remediation technologies. We demonstrate that animals can be engineered to express the microbial enzymes, organomercurial lyase (MerB) and mercuric reductase (MerA), to bioremediate methylmercury. MerA and MerB from *Escherichia coli* were functional in invertebrate (*Drosophila melanogaster*) and vertebrate (*Danio rerio*) model systems and converted methylmercury into volatile Hg^0^. The engineered animals tolerated higher exposures to methylmercury and accumulated less than half as much mercury relative to their wild-type counterparts. The outcomes of this research could be applied to reduce mercury contamination in farmed and recreationally caught fish, for species conservation, and to restore value to organic wastes contaminated with mercury.

## Introduction

Mercury is a highly toxic trace metal that is naturally occurring in the Earth’s crust. It is released into the biosphere through geogenic and anthropogenic sources(*1*). Human activities have increased the mercury concentration in the biosphere by an estimated 450% over natural levels(*1*).

The toxicity, bioavailability, and volatility of mercury is influenced by its chemical form. In the biosphere, mercury is present mainly as elemental mercury (Hg^0^), inorganic mercury (Hg^2+^), and organomercurial compounds such as methylmercury (MeHg). Hg^0^ is volatile and can be transported through the atmosphere and distributed globally(*2*). Atmospheric Hg^0^ can be oxidised through abiotic processes into Hg^2+^ and deposited onto waterbodies, soil, and plants(*1, 2*). Hg^2+^ can undergo biomethylation mediated largely by iron- and sulfur-reducing bacteria in anaerobic environments to form MeHg(*3*).

MeHg poses the greatest environmental exposure risk as it readily biomagnifies through food webs, particularly in aquatic ecosystems(*2*). The main source of mercury exposure for humans is as MeHg, typically from dietary seafood(*1, 2*). MeHg biomagnification also threatens species conservation of certain aquatic mammal, bird, and fish species(*4*). When ingested, MeHg is readily absorbed in the animal gastrointestinal tract, and MeHg assimilation typically exceeds 80%(*5*). Once in the body, MeHg is poorly excreted, and can efficiently cross the blood brain barrier, as well as the placenta(*6*). MeHg therefore acts as a potent neurotoxin, though it can also impact the cardiovascular, reproductive, and immune systems(*2, 4*). Children exposed to higher concentrations of mercury during crucial developmental stages have a lower intelligence quotient (IQ), which on a population scale can impact economic productivity(*7*).

Inorganic mercury in contrast is poorly absorbed in the animal digestive tract, and assimilation does not exceed 50%(*5, 8*). It is inefficient at crossing the blood-brain and blood-placenta barriers, and primarily accumulates in the kidneys and liver(*6*).

Many microbes natively express the enzymes MerB (organomercurial lyase) and MerA (mercuric reductase), that can detoxify MeHg in cells (Fig. 1)(*9*). Specifically, MerB catalyzes the protonolysis of MeHg to Hg^2+^, and directly transfers Hg^2+^ to MerA(*10*). MerA reduces Hg^2+^, using electrons from NADPH, to generate Hg^0^. Hg^0^ is volatilized from biomass into the atmosphere, thereby detoxifying cells of mercury.

**Fig. 1:**
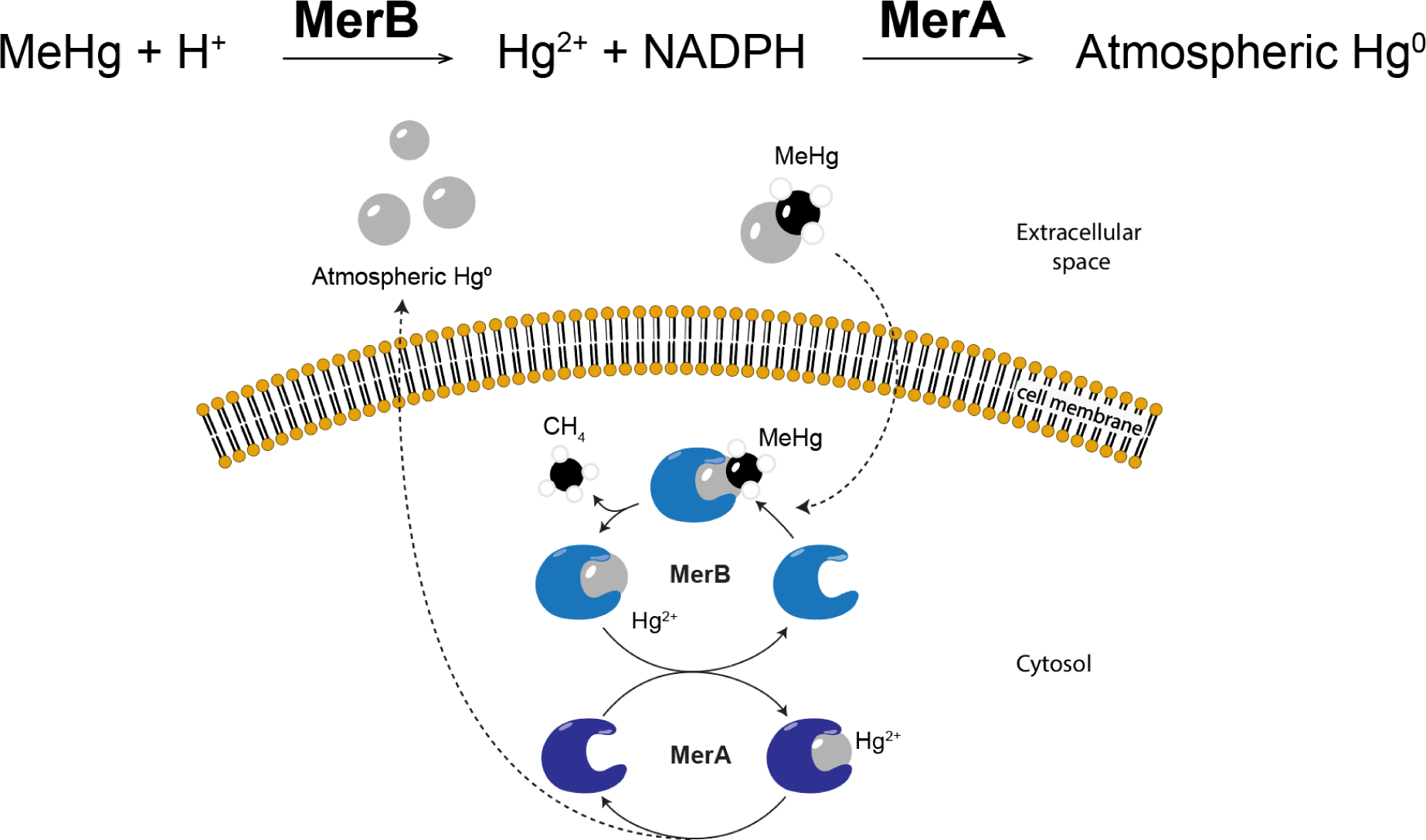
Reaction scheme for microbial MerA and MerB in cells. MerB catalyzes the protonolysis of MeHg to Hg^2+^ and directly passes Hg^2+^ to MerA. MerA reduces Hg^2+^ to volatile Hg^0^, using electrons from NADPH.

The potential of these enzymes for mercury bioremediation has been previously explored via the use of microbes or engineered plants(*11–13*). However, these cannot access mercury that is within other plants and animals, which possesses the greatest risk via dietary exposure (*9, 12–14*). Though some MeHg demethylation occurs through microbial activity in the gut microbiome(*15*), MeHg is largely absorbed in the upper gastrointestinal tract and incorporated in the animal host. Only a fraction of ingested MeHg is excreted via the biliary system and processed by the microbiome in the lower gastrointestinal tract(*16*).

Using synthetic biology, we propose that select animals could be engineered to express microbial MerA and MerB to demethylate and volatilize mercury incorporated in their biomass. This could directly disrupt MeHg biomagnification and reduce the total amount of mercury in contaminated ecosystems.

Previously, Krout *et al.,* (2022) engineered the genetic animal model, *Drosophila melanogaster* (common fruit fly) with a functional variant of MerB from the bacterial strain *Pseudomonas sp.* K-62(*17*). Interestingly, the flies engineered with only MerB typically contained 50% less total mercury in their biomass than controls. MerB expression alone can therefore contribute to mercury elimination from flies, presumably as Hg^2+^. Though this study was from the perspective of characterizing how demethylation affects MeHg toxicology and distribution, MerB expression alone is likely to be insufficient for bioremediation as Hg^2+^ would not be removed from a contaminated ecosystem. Volatile Hg^0^ in contrast, can be removed from a contaminated environment and diluted by the atmosphere. Volatilized Hg^0^ could also be trapped(*18*) in physically contained facilities, where it can be removed from the biosphere.

Engineered animals may provide unique advantages to the field of bioremediation, including access to pollutants sequestered in biomass and robust transgene biocontainment strategies. Previously we demonstrated that animals could be engineered with a high redox potential laccase from *Trametes trogii* to oxidize a variety of industrial pollutants(*19*). We now demonstrate that the genetic animal models, *D. melanogaster* (common fruit fly) and *Danio rerio* (zebrafish) can be engineered to express MerA and MerB from *Escherichia coli* to demethylate and volatilize MeHg.

## Results

### Insect mercury bioremediation

#### Engineering Fly/EcMerA+B

*MerA* and *MerB*, from *Escherichia coli* were previously shown to be functional when heterologously expressed in plants for phytoremediation(*12–14*). We reasoned they may therefore be functional when heterologously expressed in animals as they are also eukaryotes. To enable expression in flies, the genes were codon optimised for *D. melanogaster* and expressed from a truncated alpha tubulin promoter (Fig. 2A)(*20*). As the optimal expression patterning is not yet known, we selected this promoter as it provides moderate expression across a broad set of tissues. A broad expression pattern increases the likelihood that the enzymes would be expressed 1) where mercury accumulates and 2) in tissues that have favourable conditions for enzyme function. Moderate expression may also reduce their potential toxicity. The expression vectors were integrated into previously characterised landing sites using PhiC31 mediated integration: vectors containing *MerB* were integrated into a locus on Chr2, and vectors containing *MerA* were integrated into a locus on Chr3(*21*). The *mini-white* selection marker was used to select transgenic *D. melanogaster*(*22*). Flies homozygous for both *MerA* and *MerB* (Fly/EcMerA+B) were generated via breeding. A fly strain was also engineered to express the empty expression vector without *MerA* or *MerB* (Fly/Empty Vector) to serve as a control.

**Fig. 2:**
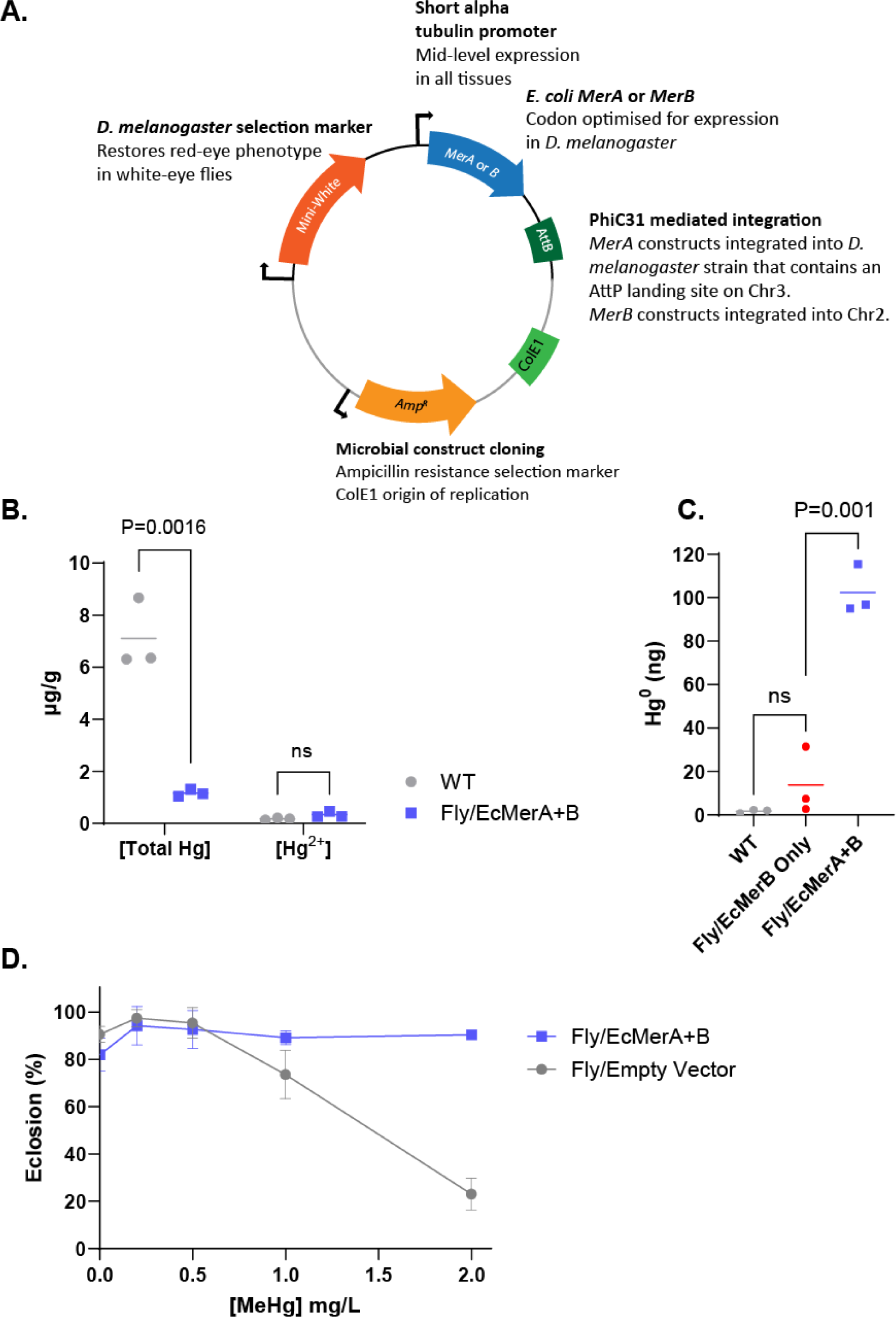
Engineering flies for mercury bioremediation. A. Diagram of the plasmids used to generate transgenic *D. melanogaster* expressing MerA and MerB. B. Total mercury and Hg^2+^ concentrations in Fly/EcMerA+B adult flies compared to WT adult flies after exposure to 0.2 mg/L MeHg during larval development. n = 3 vials with 5-10 selected adult flies each for Fly/EcMerA+B and WT. Data points are individual replicate values with horizontal bars showing the mean. The statistical analyses were conducted using unpaired t-tests, where ns = not statistically significant (p > 0.05). C. Hg^0^ volatilization after 6 days in Fly/EcMerA+B exposed to 1000 ng MeHg compared to Fly/EcMerB Only and WT flies. n = 3 vials with 150 embryos each for Fly/EcMerA+B, Fly/EcMerB Only, and WT. Data points are individual replicate values with horizontal bars showing the mean. The statistical analysis was conducted using one-way ANOVA using Dunnett’s method compared to controls, where ns = not statistically significant (p > 0.05). D. Percentage of adult fly eclosion from pupae in Fly/EcMerA+B compared to Fly/Empty Vector when exposed to 0, 0.2, 0.5, 1, and 2 mg/L MeHg from the embryo lifestage. n = 3 vials with 50 embryos each for Fly/EcMerA+B and Fly/Empty Vector. Data points are mean values ± standard deviation. Fly/EcMerA+B = *D. melanogaster* expressing *E. coli* MerA and MerB, Fly/EcMerB Only = *D. melanogaster* expressing only *E. coli* MerB, WT = wild-type *D. melanogaster*, and Fly/Empty Vector = *D. melanogaster* engineered to express the empty parent vector.

#### Fly MeHg bioconcentration assay

We assayed enzyme activity in flies co-expressing MerA and MerB by rearing larvae and controls with MeHg spiked into their diet and measuring mercury concentration in their biomass.

Triplicates of 50 WT and Fly/EcMerA+B fly embryos were added to 5 g of cornmeal diet spiked with 0.2 mg/L MeHg. The embryos were reared to maturity and the MerB breakdown product, Hg^2+^, and total mercury (organic and inorganic mercury) was measured in 5-10 adult flies within 8 h of eclosion.

Fly/EcMerA+B had 83% less total mercury than WT (Fig. 2B). Furthermore, 29% of the total mercury in Fly/EcMerA+B was as Hg^2+^, while only 2% of the total mercury in the WT flies was as Hg^2+^. This indicates that Fly/EcMerA+B is active. The lowered total mercury concentration may have been due to MeHg demethylation to Hg^2+^ and volatilization to Hg^0^, or by clearing of the MerB product, Hg^2+^, by an unknown mechanism(*17*). We therefore sought to directly measure the MerA product, volatile Hg^0^.

#### Fly Hg^0^ volatilization

We determined if MerA was active by rearing larvae and controls with MeHg spiked into their diet and measuring Hg^0^ volatilization via cold vapor atomic fluorescence spectroscopy coupled with gold amalgamation mercury preconcentration(*23*).

Triplicates of 150 embryos were added to cornmeal diet. Flies engineered to express *E. coli* MerB, without MerA (Fly/EcMerB Only) were included as a control to show whether Hg^2+^ generated by MerB can be volatilized to Hg^0^ without co-expression of MerA. After 3 days, 1,000 ng of MeHg was spiked into the diet and the sample vessels were immediately connected to an apparatus which captured volatilized Hg^0^ onto gold coated quartz sand Hg^0^ traps for six days.

Substantial Hg^0^ was detected in the Fly/EcMerA+B strain treatment with only trace amounts of Hg^0^ detected in the WT strain treatment (Fig. 2C). Fly/EcMerB Only volatilized slightly more Hg^0^ than WT, though it was not statistically significant. The larvae engineered to express MerA and MerB volatilized 10% of the MeHg added to their diet into Hg^0^ over the course of the 6-day assay. These results confirmed that both MerA and MerB are active in flies and capable of converting MeHg to Hg^2+^, which is subsequently reduced and volatilized as Hg^0^.

**Fig. S1:**
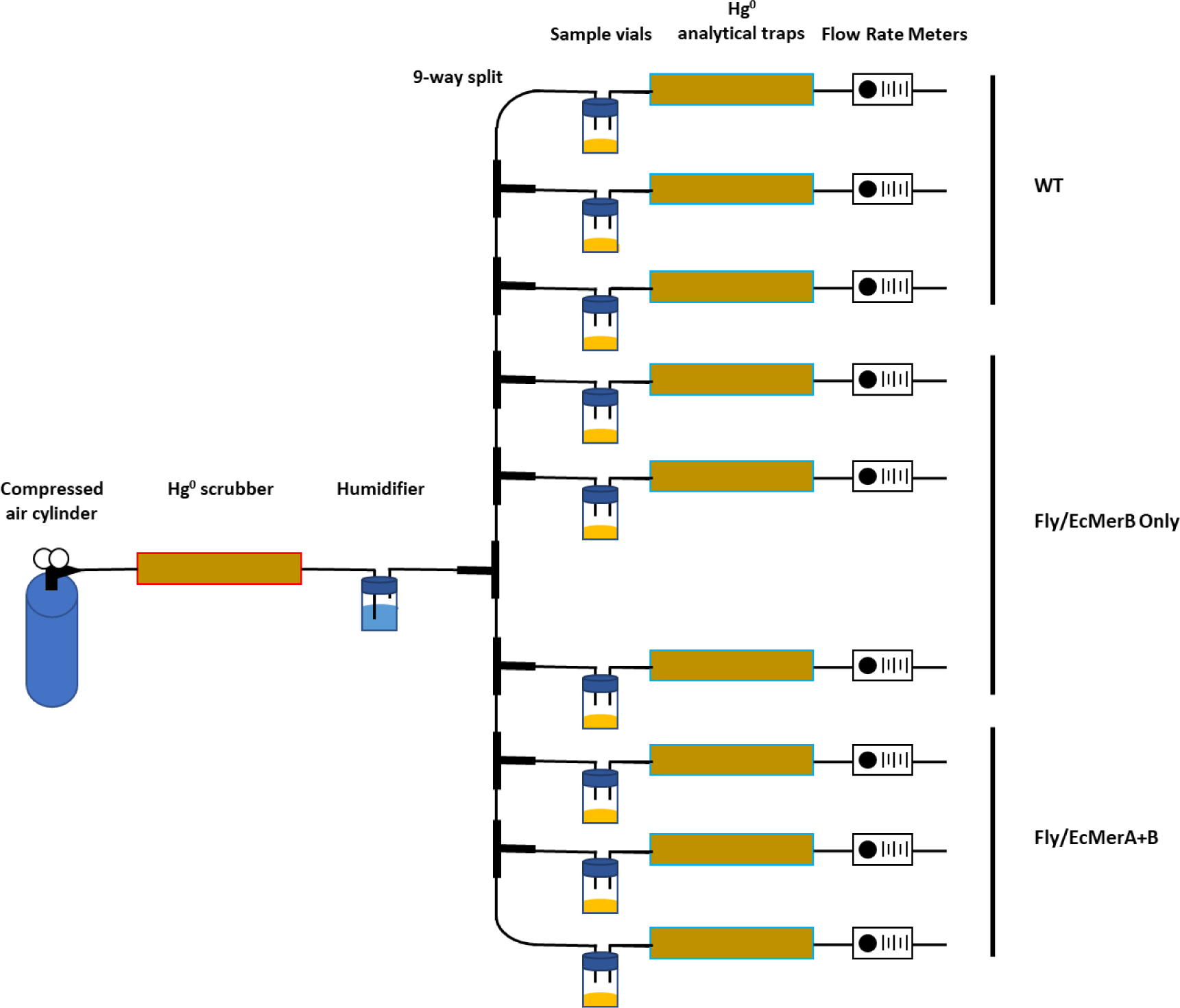
schematic diagram of the Hg^0^ evolution and trapping apparatus used in the fly experiments.

#### Fly toxicity assay

Animals engineered for bioremediation will be exposed to the toxic effects of the pollutants they are targeted to remediate. Understanding how they may be impacted is important to ensure their viability. For instance, Krout *et al.,* (2022) showed that MeHg significantly reduced eclosion in WT *D. melanogaster* and expression of MerB in *D. melanogaster* could restore eclosion(*17*). Other development stages such as hatching, pupariation, and longevity are more subtly affected by MeHg exposure(*24*). We therefore tested whether Fly/EcMerA+B is also resistant to the toxic effects of mercury.

We assayed eclosion in Fly/EcMerA+B compared to controls by adding 50 fly embryos to cornmeal diet spiked with 0, 0.2, 0.5, 1, and 2 mg/L MeHg. Fly/Empty Vector was used as a control to account for potential genetic background fitness defects. The embryos developed to adulthood, which takes approximately 10 days.

Fly/EcMerA+B eclosion was not affected by MeHg across the tested MeHg concentrations (Fig. 2D). Meanwhile, only approximately 20% of the Fly/Empty Vector controls eclosed when exposed to 2 mg/L MeHg. This result demonstrates that expression of MerA and MerB confers resistance to MeHg exposure toxicity.

### Fish mercury bioremediation

#### Engineering Fish/EcMerA+B

To explore mercury bioremediation in vertebrates, *E. coli* MerA and MerB were heterologously expressed in zebrafish (*Danio rerio*) (Fig. 3A).

**Fig. 3:**
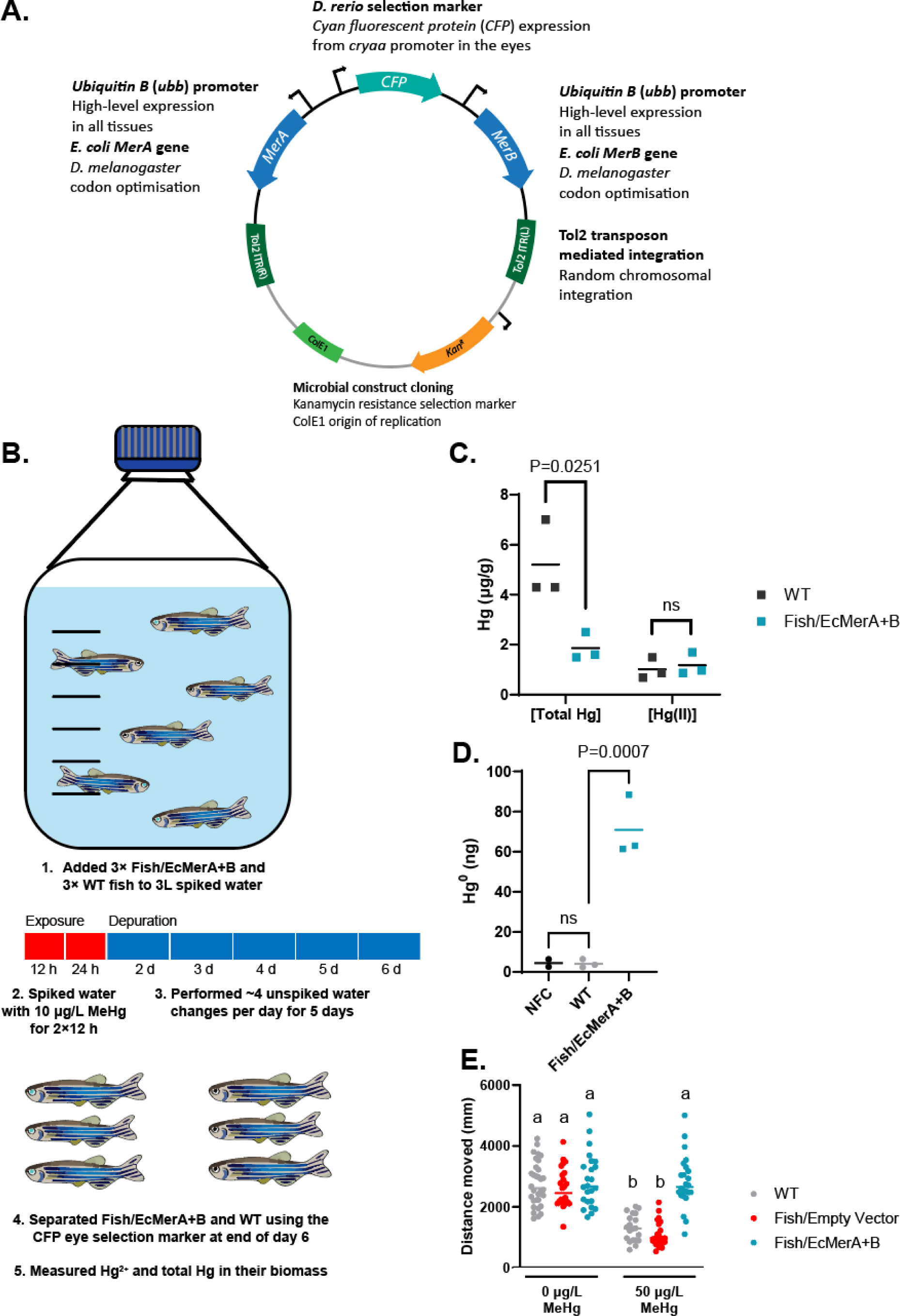
Engineering fish for mercury bioremediation. A. Diagram of the plasmid used to generate transgenic *D. rerio* expressing MerA and MerB. B. Adult zebrafish mercury bioconcentration assay design overview. C. Total mercury and Hg^2+^ concentrations in adult Fish/EcMerA+B compared to WT. Fish were exposed to 10 μg/L MeHg for 2×12 h, before depuration with unspiked water for 5 days. n = 3 fish each for Fish/EcMerA+B and WT. Data points are individual replicate values with horizontal bars showing the mean. The statistical analyses were conducted using unpaired t-tests, where ns = not statistically significant (p > 0.05). D. Hg^0^ volatilization after 7 days in Fish/EcMerA+B larvae exposed to 1000 ng MeHg compared to WT and NFC. n = 3 bottles with 200 larvae each for Fish/EcMerA+B and WT. n = 2 bottles with no larvae for NFC. Data points are individual replicate values with horizontal bars showing the mean. The statistical analysis was conducted using one-way ANOVA using Dunnett’s method, where ns = not statistically significant (p > 0.05). E. Swimming fitness in Fish/EcMerA+B larvae exposed to 0 μg/L and 50 μg/L MeHg compared to WT and Fish/Empty Vector. n = at least 21 larvae each for Fish/EcMerA+B, Fish/Empty Vector, and WT. Data points are individual replicate values with horizontal bars showing the mean. The statistical analysis was conducted using one-way ANOVA using Dunnett’s method, where treatments with the same letter are not statistically significantly different from each other (p > 0.05), and treatments with different letters are statistically significantly different from each other (p < 0.05). Fish/EcMerA+B = *D. rerio* expressing *E. coli MerA* and *MerB*, WT = wild-type *D. rerio*, NFC = No Fish Control.

As MerA and MerB were well tolerated when co-expressed in flies, we assembled a dual expression vector to simplify establishing strains. *E. coli MerA* and *MerB* that were codon optimised for *D. melanogaster* were both expressed from the zebrafish *ubiquitin B* promoter (ubb) for high-level expression across a broad set of tissues (Fish/EcMerA+B). Following embryo microinjection, plasmids were randomly integrated into the zebrafish genome via Tol2 transposase mediated integration. Larvae were screened for cyan fluorescent protein (CFP) in the eyes and transgenic lines were established. The fish were also engineered to express the empty dual expression vector without MerA or MerB (Fish/Empty Vector) to serve as a control.

#### Adult zebrafish mercury bioconcentration assay

We first evaluated the performance of Fish/EcMerA+B by exposing adults to MeHg and determining if they were capable of clearing it from their biomass.

Fish/EcMerA+B adults (3 five-month-old adults; F2 generation heterozygotes) and WT adults (3 five-month-old adults) were added to 3 L water in the same glass Duran™ bottle (Fig. 3B). During the initial exposure stage, the water was changed twice in 12 h with water containing 10 μg/L MeHg. After 24 hours, the depuration stage began and 3 L of clean water was changed approximately 4 times daily (depending on ammonia concentration measured in the water) until day 6. At the end of day 6, all the fish were separated using the CFP eye selection marker. Total mercury and Hg^2+^ concentrations were measured in their biomass.

After 6 days, the WT zebrafish tissue total mercury concentration was 5.2 μg/g on average (Fig. 3C), similar to the highest mercury concentrations in wild caught sharks(*25*). Fish/EcMerA+B had 1.9 μg/g total mercury on average, which was 64% lower than WT. Though the Fish/EcMerA+B Hg^2+^ concentration was not statistically significantly different compared to WT, the majority of the mercury in Fish/EcMerA+B was determined to be Hg^2+^.

#### Zebrafish larvae Hg^0^ assay

We determined if MeHg was removed from Fish/EcMerA+B as Hg^0^ by measuring Hg^0^ volatilization via cold vapor atomic fluorescence spectroscopy coupled with gold amalgamation mercury preconcentration.

Fish/EcMerA+B larvae (200 larvae at 4 days post fertilisation (dpf); F2 generation heterozygotes), WT larvae (200 larvae at 4 dpf), and No Fish Control (NFC) were added to 200 mL E3 media containing 5 μg/L MeHg (1000 ng total mercury). Sample vessels were promptly connected to the Hg^0^ apparatus. After 24 hours, the water was exchanged with 200 mL clean E3 media daily for a total of 7 days.

Substantial Hg^0^ was detected from Fish/EcMerA+B (Fig. 3D). Only trace amounts of Hg^0^ were detected from the WT strain. These results confirm that both MerA and MerB are active in fish and capable of converting MeHg to Hg^0^. The larvae engineered to express MerA and MerB volatilized 7% of the MeHg added to their water during the exposure stage into Hg^0^ over the course of the 7-day assay.

#### Zebrafish swimming fitness assay

We measured whether Fish/EcMerA+B are resistant to the toxic effects of MeHg by assessing swimming fitness.

Fish/EcMerA+B larvae and controls were exposed to 0 μg/L and 50 μg/L MeHg for 24 h. Larvae were then exposed to 6 minutes in the light, 6 minutes in the dark, repeated twice. Fish/Empty Vector was used as a control to account for potential genetic background fitness defects.

Fish/EcMerA+B swimming distance was not affected by 50 μg/L MeHg exposure compared to 0 μg/L MeHg (Fig. 3E). Meanwhile, both WT and Fish/Empty Vector controls swam approximately half the total distance when exposed to 50 μg/L MeHg, compared to 0 μg/L MeHg.

## Discussion

MeHg that has biomagnified in food webs causes significant environmental harm and has previously been inaccessible to remediation technologies, such as microbial bioremediation and phytoremediation. We demonstrated for the first time that animals can remediate MeHg by converting it to volatile Hg^0^. We engineered *D. melanogaster* and *D. rerio* to express MerA and MerB from *E. coli*. The transgenic flies and zebrafish had 83% and 64% less total mercury in their biomass than their WT counterparts, respectively. Furthermore, compared to WT, a higher proportion of the remaining mercury in their biomass was in the form of Hg^2+^, which does not bioaccumulate as readily as MeHg(*5, 8*). Animals engineered for mercury bioremediation will be able to tolerate greater exposure to mercury than WT, while potentially reducing mercury levels to other wildlife.

Reducing Hg^2+^ to volatile Hg^0^ and expanding these metabolic capabilities to vertebrates demonstrated in the present study enables applications that aim to reduce the total mercury content of a polluted site, facilitate the extraction and collection of mercury from contaminated wastes, or protect higher trophic levels from mercury toxicity.

Specific examples include engineering farmed or recreational fish stocks. Consumers could gain all the nutritional benefits of fish(*26, 27*), with reduced exposure to mercury. It could also be applied for species conservation, where aquatic invertebrates or fish at lower trophic levels that may be responsible for the majority of mercury biomagnification, could be engineered to remediate ecosystems. This could also reduce the mercury concentration in recreationally or commercially harvested fish. The release of transgenic organisms is likely to be controversial. However, a poll of Australians residing near polluted waterways indicated greater support for the use of transgenic fish for bioremediation than conventional remediation technologies(*28*).

Using animals for mercury bioremediation could also be implemented in physically contained systems. Insects (e.g. black soldier flies) can be used to process organic wastes with high concentrations of mercury, such as municipal biosolids and offal from fisheries waste (1, 29). Municipal biosolids are not currently decontaminated and a quarter of the mercury inventory released to the environment is from municipal sewage (1). Transgenic insects reared on biosolids may provide novel means of decontamination. Mercury can be removed as Hg0 from both the pupae and the spent organic waste (frass), which can be used in other applications such as animal feed or fertilizer, respectively.

Effective transgene biocontainment is critical for any bioremediation approach that uses engineered organisms, yet is a major challenge for plants and microbes(*30, 31*). Transgene biocontainment is already well established in animals. Farmed fish are currently sterilized by inducing triploidy or surgical sterilization(*32*). Biocontainment in insects has been demonstrated in several ways including: engineered reproductive barriers that prevent interbreeding with their WT counterparts(*33*), conditional lethality(*34*), as well as gene knockouts resulting in flightless insects that could be reared in a contained facility, but would impose a significant fitness cost in case of accidental release(*35*).

The fate of the Hg^0^ produced by transgenic animals would depend on the application. Hg^0^ released by animals reared in physically contained facilities, such as farmed fish and insects, could be trapped(*18*) and safely removed from the biosphere. For animals engineered for bioremediation in the environment, volatile Hg^0^ would be diluted in the atmosphere. This may be acceptable in circumstances where the MeHg burden is high and the risk of harm to people or animals would be substantially reduced by conversion to volatile Hg^0^. Understanding the routes of mercury biomagnification and exposure in a given ecosystem will be necessary to ensure implementation results in the highest benefit.

## Acknowledgments

We are grateful to the zebrafish facility technical staff at Macquarie University for caring for our zebrafish, specific thanks to Jason Martin-Powell, Kade Smith, and Cheryl Song for their contributions to animal husbandry and their expertise in animal care. The authors thank Aidan J Peterson from the University of Minnesota for providing the *D. melanogaster* balancer strains.

## Ethics

Experimental protocols were approved by Macquarie University Animal Ethics (ARA 2021/025-2) and Institutional Biosafety Committees (#7801).

## Funding

International Technology Center Pacific (ITC-PAC) under Contract No. FA520920P0100 (MASELKO).

CSIRO Environment (SA, JK).

## Author contributions

Conceptualization: KT, MASELKO

Methodology: KT, JK, PMC, CP, MORSCH, SA, MASELKO

Investigation: KT, JK, PMC, CP Visualization: KT

Funding acquisition: SA, MASELKO

Project administration: KT, JK, PMC

Supervision: CP, MORSCH, SA, MASELKO

Writing – original draft: KT

Writing – review & editing: KT, JK, PMC, CP, MORSCH, SA, MASELKO

## Competing interests

This research was filed under patent application WO/2023/183984.

## Data and materials availability

All data are available in the main text or the supplementary materials.

## Materials and Methods

### Preparation of Hg stocks

All glass and plasticware used in analytical procedures were cleaned prior to use by soaking in 10% v/v hydrochloric acid for at least 24 hours, then rinsed with copious amounts of ultra-pure water, then allowed to dry in a laminar flow hood. ‘Ultrapure’ reverse osmosis water (18 MΩ.cm, Millipore, low in mercury) was used to prepare all solutions. Total mercury and Hg^2+^ standards were prepared from certified stock solutions (Accustandard, USA). MeHg standards were prepared from methylmercury chloride (Sigma-Aldrich, USA) and verified as per US EPA 1630 (*36*) against certified total mercury standards. As per US EPA 1630(32), all MeHg quantities and concentrations are reported as MeHg as Hg: the quantity of Hg, that is within the MeHg compound. This facilitates comparisons and mass balance conversions between MeHg (as Hg) and Hg^2+^, total Hg, and Hg^0^.

### Fly methods

#### Fly expression plasmid assembly

The *E. coli MerA* and *MerB* enzyme sequences (Genbank IDs: AAB59078 and AAB49639, respectively) were codon optimised for expression in *D. melanogaster* using the Integrated DNA Technologies (IDT) codon optimisation tool. The *D. melanogaster* Kozak consensus sequence: AATCTTACAAA was added immediately preceding the start codon. At least 20 bp of homology arms to the destination plasmid were added upstream and downstream of these sequences. These sequences were ordered as gBlocks from IDT and assembled (NEBuilder® HiFi DNA Assembly Master Mix (New England Biolabs (NEB))) into the pMC-1-1-1(*19*) expression plasmid linearised with NotI-HF. The plasmids were verified by Sanger Sequencing. The plasmid DNA sequences are listed in Supplementary Table S4.

#### Fly strains, transgenesis, and husbandry

Canton S wild-type flies were sourced from the Bloomington Drosophila Stock Centre (RRID: BDSC #64349). Balancer fly strains SGSB (w^−^; sp1 / CyO-GFP, mW^+^; Sb / TM6b,Tb,Hu,e) and *ST (w^−^; Pin / Cyo*; I(3),e / TM6C,Sb,Tb,e) were kind gifts from Aidan J Peterson from the University of Minnesota.

Enzyme expression plasmids were sent to BestGene Inc (Chino Hills, Ca) for plasmid DNA midipreps, microinjections facilitated by φC31 mediated genomic integration, and outcrossing of transgenic lines to balancer strains. Double homozygous lines (Fly/EcMerA+B) were generated by crossing *D. melanogaster* engineered to express MerB (Fly/EcMerB Only) with *D. melanogaster* engineered to express MerA (Fly/EcMerA Only) using double balancer strains.

Flies were maintained on cornmeal diet based on the Bloomington Drosophila Stock Centre (BDSC) standard Nutri-Fly formulation (catalogue number 66-113; Genesee Scientific). Flies were reared in a controlled environment room at 25°C, 75% humidity, and a 12 h light-dark cycle with a 30 min transition period. Biosafety approval was granted by the Macquarie University Institutional Biosafety Committee (#7801).

#### Fly MeHg bioconcentration assay

Fifty grams of 55-65 *°*C cornmeal diet was spiked with 0.2 mg/L MeHg in a 150 mL glass Duran™ bottle. Aliquots of 5 g spiked cornmeal diet were then set into 6 clean 40 mL glass vials (Shimadzu, 226-50581-00).

Fly embryos (<12 h old) were harvested from grape-agar medium plates seeded by a mating population of 150-300 adult flies. Triplicates of 50 embryos for WT and Fly/EcMerA+B were added to the spiked cornmeal diet and the offspring larvae were reared until completion of their lifecycle. Hg^2+^ and total mercury (organic and inorganic mercury) concentrations were measured in 5-10 adult flies within 8 h of eclosion from each sample.

#### Fly Hg^0^ volatilization

Embryos (<12 h old) were harvested from grape-agar medium plates seeded by a mating population of 150-300 adult flies. Triplicates of 150 embryos for WT, Fly/EcMerB Only, and Fly/EcMerA+B were added to 5 g of cornmeal diet set into 150 mL glass Duran™ bottles. After 3 days, 1,000 ng of MeHg was spiked into the diet and the sample vessels were immediately connected to a purpose-built apparatus that was developed to measure Hg^0^ released by the study organisms (the Hg^0^ apparatus). Prior to connecting the sample vessels, the Hg^0^ apparatus was pre-purged for 10 min with Hg^0^ scrubbed flowing air.

A diagram of the Hg^0^ apparatus is shown in Fig. S1. Instrument grade air cylinders (BOC, Gas Code 054) fitted with a dual stage regulator (Coregas, 600215, FM53-300/1.5) supplied constant air flow into the apparatus at approximately 0.05 L/min. Silicone tubing piped air into a gold-coated glass bead Hg^0^ quartz sand scrubber (Brooks Rand, SKU 03034), connected by Teflon adapters (Brooks Rand, SKU 08401). The Hg^0^ scrubbed air was humidified by piping Teflon tubing (Brooks Rand, SKU 08406) directly into ∼100 mL milli-Q water inside a 150 mL glass Duran™ bottle fitted with a GL45 lid with 2 GL14 ports (Duran™, 1129750). The Teflon tubing was then split 9 ways using perfluoroalkoxy (PFA) 3-way connectors (Swagelok, PFA-220-3), and connected to the sample vials described above with a 2-port GL45 lid. Any volatilized Hg^0^ from the samples flowed into the headspace, and amalgamated onto downstream Hg^0^ analytical gold coated quartz sand traps (Brooks Rand, SKU 08401). The samples were connected to Hg^0^ analytical traps via Teflon tubing and 2-way connectors. Flow rate meters (Dwyer Instruments, VFA-21-SSV) were attached at the end and used to maintain consistent flow rate between the samples.

The Hg^0^ apparatus was run for 6 days. Following the experiment, the traps were disconnected, sealed with Teflon plugs and stored at room temperature before analysis.

#### Fly toxicity assay

Aliquots of 50 g of 55-65 *°*C cornmeal diet were spiked with 0, 0.2, 0.5, 1, or 2 mg/L MeHg in 150 mL glass Duran™ bottles. Aliquots of 5 g spiked cornmeal diet were then set into clean 40 mL glass vials (Shimadzu, 226-50581-00).

Fly embryos (<12 h old) were harvested from grape-agar medium plates seeded by a mating population of 150-300 adult flies. Triplicates of 50 embryos for WT and Fly/EcMerA+B were added to each spiked cornmeal diet concentration and incubated for 20 days. The total number of developed pupae and the total number of eclosed adults were then counted.

### Fish Methods

#### Zebrafish expression plasmid assembly

pSC1-2 was used to generate the zebrafish enzyme expression plasmids. pSC1-2 contains a Zebrafish *ubb* promoter upstream of a NotI restriction site and a rabbit HBB2 polyA signal sequence. It contains a *cryaa* eye promoter facing in the opposite direction to *ubb*, driving the *mCerulean* (*CFP*) selection marker with a bGH polyA signal sequence. Flanking these are the Tol2 Inverted Terminal Repeat (ITR) left and right border recognition sequences.

NEBuilder HiFi DNA Assembly Master Mix (NEB) was used for all assembly reactions. Q5® High-Fidelity 2X Master Mix (NEB) was used for all polymerase chain reactions (PCRs).

A second *ubb* promoter with an AscI restriction site in between an additional rabbit HBB2 polyA signal sequence was installed between the pSC1-2 bGH polyA signal sequence and the Tol2 ITR left border. This was achieved by 3-part assembly of a PCR amplicon consisting of the zebrafish *ubb* promoter with a downstream AscI restriction site, a second PCR amplicon containing the AscI restriction site upstream of the rabbit HBB2 polyA signal sequence, and pSC1-2 cut with BbsI. This plasmid, pKT1-2.Ubb::Empty.Ubb::Empty, was used as the empty parent vector.

The *MerA* open reading frame (ORF) was PCR amplified from pKT111.EcMerA and assembled into pKT1-2.Ubb::Empty.Ubb::Empty cut with NotI. The *MerB* ORF was PCR amplified from pKT111.EcMerB and assembled into pKT1-2_Ubb::EcMerA.Ubb::Empty cut with AscI. The Kozak sequence GCCACC was added upstream to both the *MerA* and *MerB* ORFs. This plasmid, pKT1-2.Ubb::EcMerA.Ubb::EcMerB, was used to generate Fish/EcMerA+B.

All plasmids were sequence verified by Plasmidsaurus whole plasmid sequencing. The plasmid DNA sequences, pKT1-2.Ubb::Empty.Ubb::Empty and pKT1-2.Ubb::EcMerA.Ubb::EcMerB, are listed in Supplementary Table S4.

#### Zebrafish strains, transgenesis, and husbandry

Wild-type zebrafish (AB/Tübingen background) were used for experimental controls and as the genetic background for microinjections.

Fish/EcMerA+B and Fish/Empty Vector were generated by Tol2-based transposon mediated random genomic integration methods(*37*). Midiprepped plasmid DNA (25 ng/μL), Tol2 transposase mRNA (25 ng/μL), and 0.1% phenol red was microinjected into one cell stage WT zebrafish embryos as described previously(*38, 39*). Transgenic fish were screened at 2-4 dpf for CFP expression in the eyes using a CFP filter (Leica, 10447409) on a fluorescence stereomicroscope (Leica, M205 FCA).

Zebrafish (*Danio rerio*) were maintained in standard conditions(*40, 41*). All water and E3 media in zebrafish husbandry were prepared as described previously(*40*). Experiments were approved by the Macquarie University Animal Ethics Committee (ARA 2021/025-2).

#### Adult zebrafish MeHg bioconcentration assay

Triplicates of five-month-old zebrafish adults for WT and Fish/EcMerA+B were added to 3 L system water into a 3 L glass Duran™ bottle. During the initial MeHg exposure stage, the water was changed twice in 12 h with system water spiked with 10 μg/L MeHg (24 h total MeHg exposure). After 24 hours, the depuration stage began, and 3 L of unspiked system water was changed approximately 4 times daily (depending on ammonia concentration measured in the water) until day 6. Ammonia concentration was measured using the ammonia (NH_3_, NH ^+^) test kit (API, LR8600) and the water was changed if concentrations exceeded 0.25-0.5 mg/L. At the end of day 6, all the fish were euthanized and separated using the CFP eye selection marker. Total mercury and Hg^2+^ concentrations were measured in their biomass. Also see Fig. 3B for a diagram of the experimental procedure.

#### Zebrafish larvae Hg^0^ volatilization assay

Triplicates of 200 4-dpf zebrafish larvae for WT and Fish/EcMerA+B were added to 200 mL E3 media spiked with 5 μg/L MeHg (1000 ng total Hg) in 500 mL glass Duran™ bottles. Samples bottles were connected within 10 minutes to the Hg^0^ apparatus that had been pre-purged for 10 minutes with Hg^0^ scrubbed flowing air. After 24 hours, the water was exchanged with 200 mL unspiked E3 media daily. The Hg^0^ apparatus was kept connected for a duration of 7 days.

The fish Hg^0^ apparatus in the zebrafish experiments (Fig. S2) was the same as the fly Hg^0^ apparatus (“Fly Hg^0^ volatilization” methods section, (Fig. S1)) with the following exceptions. An aquarium air pump (Aqua One, Precision 2500) was used to supply constant air flow instead of instrument grade air cylinders. The Hg^0^ scrubber was therefore replaced daily. The air was not humidified prior to the sample vials. The tubing was split 8-ways instead of 9-ways to maintain the air flow rate at approximately 0.05 L/min.

**Fig. S2:**
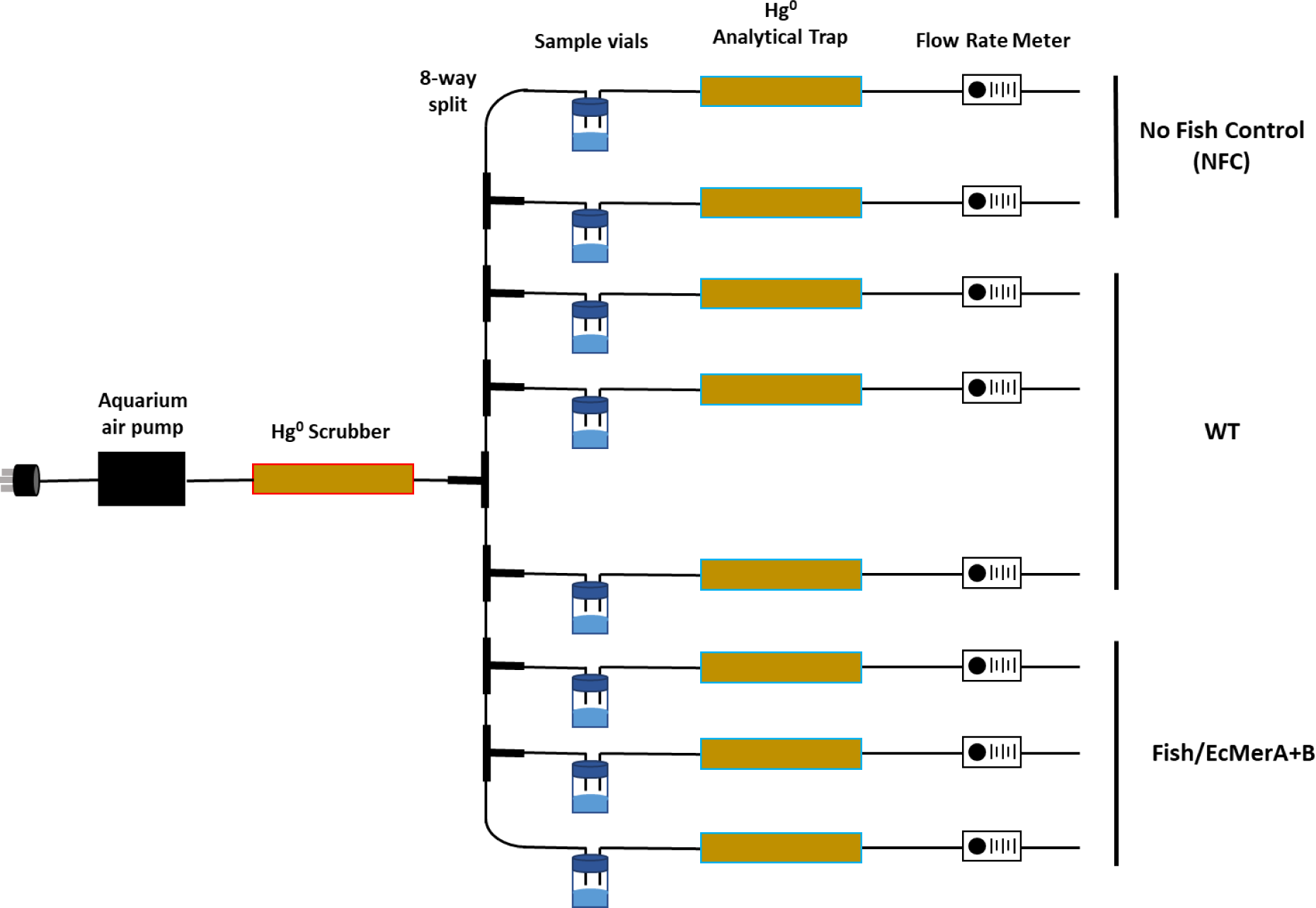
schematic diagram of the Hg^0^ evolution and trapping apparatus used in the fish experiments.

#### Zebrafish larvae swimming assessment

Larval swimming assessment in response to light-dark cycles was performed within a Zebrabox (Viewpoint) automated zebrafish movement recording device with ZebraLab (Viewpoint) software according to previously established protocols(*42*).

Larvae were incubated at 28°C in a 13:11 light-dark photoperiod from 0-dpf. Duplicates of 20 4-dpf zebrafish larvae for WT, Fish/Empty Vector, and Fish/EcMerA+B were added to 10 mL E3 media spiked with 0 and 50 μg/L MeHg in 40 mL glass vials. After 24 h, the water was exchanged with unspiked E3 media. At 5-dpf, at least 8 zebrafish larvae per sample were moved into 48-well plates, containing 1 zebrafish larva per well in 500 μL E3 media.

The 48-well plates were moved to the Zebrabox and acclimatized for 1 hour in light conditions. Then the animals were exposed to 6 min light, and 6 min in the dark, repeated twice. The total distance travelled by each animal during the 24-min test period was calculated.

### Analysis of Hg species in fly and zebrafish biomass

#### Biological tissue sample digestion

Digestion procedures were based upon the methods described by Carrasco & Vassileva (2014)(*43*) and Ramalhosa *et al.,* (2001)(*44*). Known masses of tissue (up to 0.1 g) was weighed into Teflon digestion vessels. To each vessel, 6 mL of 25% m/m potassium hydroxide (Sigma, USA) in methanol (Sigma, HPLC grade, USA) was added. The solutions were thoroughly mixed, then heated in a microwave digester (MARS Xpress), ramping to 70°C in 3 min, then holding at 70°C for 8 min. After cooling, samples were centrifuged for 20 min at 3,000 rpm to sediment out any remaining solids.

#### Total mercury analysis

Total mercury concentrations were quantified by cold vapor generation atomic fluorescence spectrometry (CV-AFS) with single-stage gold amalgamation(*23*). A 0.1 mL aliquot of digest solution was added to 80 mL of ultrapure water in a glass reaction vessel, fortified to contain 0.5% v/v bromine monochloride. The mixture was allowed to stand for at least 3 hours, after which 0.1 mL of 20% w/v hydroxylamine hydrochloride reagent was added (reaction time >3 min), followed by the addition of 0.5 mL of 20% w/v tin chloride dihydrate solution. The vessel was immediately sealed, connected to a purpose-built purge-trap apparatus and the solution purged with mercury-free nitrogen (Coregas, Australia) for 20 min. The evolved mercury was trapped on a gold coated glass bead trap (Brooks Rand, USA). The trap was then removed and connected to a thermal desorption rig connected to the AFS detector (Brooks Rand Model III, USA). The trap was purged with high purity, mercury-free argon gas (Coregas, Australia) and the coil heated (450-500°C) to release the mercury in the gas stream and then into the AFS detector.

#### MeHg analysis

MeHg concentrations were determined by aqueous phase ethylation and gas chromatography-atomic fluorescence spectrometry(*23*). A 0.03 mL aliquot of digest solution was added to 100 mL of ultrapure water in a glass reaction vessel and buffered to pH 4.9±0.1 by addition of 2 M acetate buffer solution. To this, 0.1 mL of 2% w/v sodium tetraethyl borate in 1% w/v potassium hydroxide solution was added. The vessel was immediately sealed and solution allowed to react for 15 min, before being purged with mercury-free nitrogen (Coregas, Australia) for 20 min, collecting the evolved ethylmethlymercury on a Tenax trap (Brooks Rand, USA). The trap was then removed and connected to a thermal desorption unit). The heating coil was heated to 450-500°C for 30 s causing the alkyl mercury compounds to be desorbed into a stream of high purity, mercury-free argon gas. The mercury compounds were separated by gas chromatography (Brooks Rand GC module, Chromasorb W, heated to 36°C), followed by reduction to Hg^0^ in an online pyrolysis tube (Brooks Rand, USA), and then detection by AFS (Brooks Rand Model III, USA).

#### Hg^2+^ analysis

Hg^2+^ was determined by selective evolution of Hg^0^ followed by gold amalgam preconcentration and CVAFS. A 0.1 mL aliquot of digest solution was added to 80 mL of ultrapure water in a glass reaction vessel, fortified to contain 0.5% v/v hydrochloric acid (Merck Tracepur). To this, 0.5 mL of 20% w/v tin chloride dihydrate was added and the vessel immediately sealed and solution purged with mercury-free nitrogen (Coregas, Australia) for 20 min. The evolved Hg^0^ was preconcentrated on gold coated glass bead traps (Brooks Rand, USA). The trap was then removed and placed into a stream of high purity, mercury-free argon gas (Coregas, Australia) and the trap heated to 450-500°C to release the mercury, which was then detected by CVFS (Brooks Rand Model III, USA).

#### Quality Control/Quality Assurance

Atomic Fluorescence Spectrometers were calibrated daily on use. Validation of the digest and analysis methods involved the analysis of reference materials with certified concentrations of both total and MeHg, along with pre-digestion spikes as a check to ensure the digestion and analysis procedures did not result in inter-conversion of mercury species. Reference materials included DOLT-5 (fish protein, National Research Council, Canada) and SRM2976 (mussel tissue, National Institute of Standards & Technology, USA). Results for the reference materials are shown in the table below.

| Sample |  | Total Hg (ng/g) | MeHg (ng/g) |
| --- | --- | --- | --- |
| DOLT-5 | Measured* | 442 ± 8 | 110 ± 19 |
|  | Certified | 440 ± 180 | 119 ± 58 |
| SRM2976 | Measured* | 60.3 ± 4.9 | 31.7 ± 1.8 |
|  | Certified | 61.0 ± 3.6 | 28.09 ± 0.31 |

*\*Measured values are the mean of 4 replicates ± 1 standard deviation*

Pre-digestion sample spikes of MeHg and Hg^2+^ were also used to ensure digestion and analysis did not alter mercury speciation (mean recoveries of 105% and 99% for MeHg and Hg^2+^, respectively). Analytical blanks and recovery spikes were also included in all sample batches. Recoveries of MeHg, Hg^2+^ and total mercury spikes were 90-111%. Absolute blank values were typically less than 15% of the concentrations measured in the samples. All reported data has been corrected for the blank values.

#### Measurement of evolved Hg^0^ from the Hg^0^ apparatus

In the analytical laboratory the Hg^0^ analytical traps were connected to a thermal desorption unit and purged with a stream of high purity, mercury-free argon gas (Coregas, Australia). The trap was heated to 450-500°C and the released mercury detected by AFS (Brooks Rand Model III, USA).

#### Statistics

The P values, sample numbers, and statistical tests used are stated in the figure legends. Statistical significance level for the analyses were <0.05.

#### Figures

Figures were made using GraphPad Prism 10, Biorender, and Adobe Illustrator.

**Supplementary Table S1:** *D. melanogaster* strains.

| Strain Description | BDSC # | Genotype | Reference |
| --- | --- | --- | --- |
| WT | 64349 | Canton S | (45) |
| Chr2 AttP3 for microinjections | 9752 | y<sup>1</sup> w<sup>1118</sup>; PBac{y[+]-attP-3B}VK00037 | (46) |
| Chr3 AttP3 for microinjections | 9748 | y<sup>1</sup> w<sup>1118</sup>; PBac{y+ -attP-3B}VK00031 | (46) |
| SGSB Chr2 and Chr3 balancer | N/A | w<sup>1118</sup>; sp1 / CyO-GFP, mW<sup>+</sup>; Sb / TM6b,Tb,Hu,e | Aidan J Peterson (UMN) |
| *ST Chr2 and Chr3 balancer | N/A | w<sup>1118</sup>; Pin / Cyo*; l(3),e / TM6C,Sb,Tb,e | Aidan J Peterson (UMN) |
| Fly/EV | N/A | y<sup>1</sup> w<sup>1118</sup>; PBac{pMC1-1-1}VK00037 | This study |
| Fly/EcMerB | N/A | y<sup>1</sup> w<sup>1118</sup>; PBac{pKT111.EcMerB}VK00037 | This study |
| Fly/EcMerA | N/A | y<sup>1</sup> w<sup>1118</sup>; PBac{pKT111.EcMerA}VK00031 | This Study |
| Fly/EcMerA+B | N/A | y<sup>1</sup> w<sup>1118</sup>; PBac{pKT111.EcMerB}VK00037/<br>PBac{pKT111.EcMerB}VK00037;<br>PBac{pKT111.EcMerA}VK00031/<br>PBac{pKT111.EcMerA}VK00031 | This Study |

**Supplementary Table S2:**
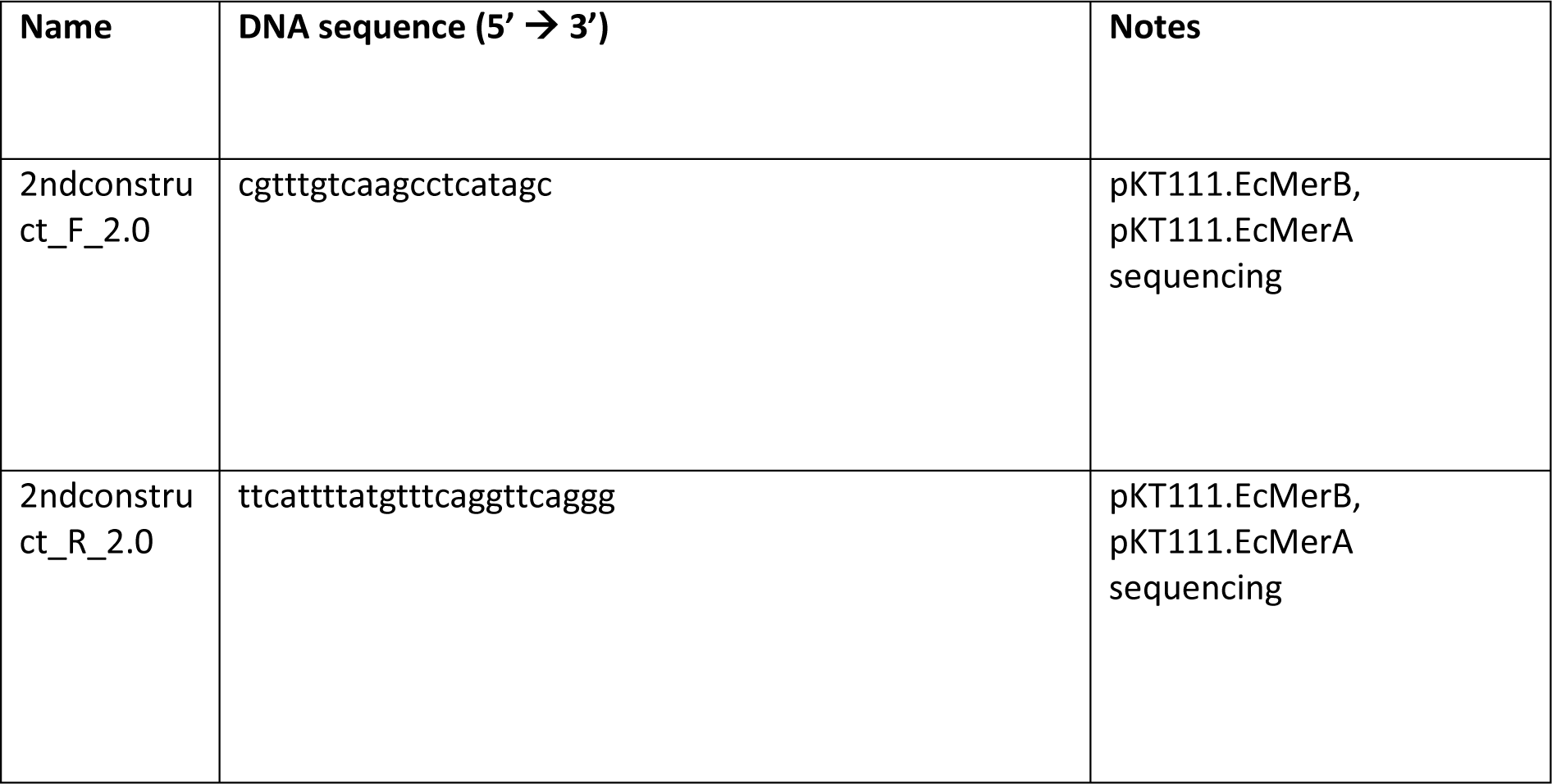
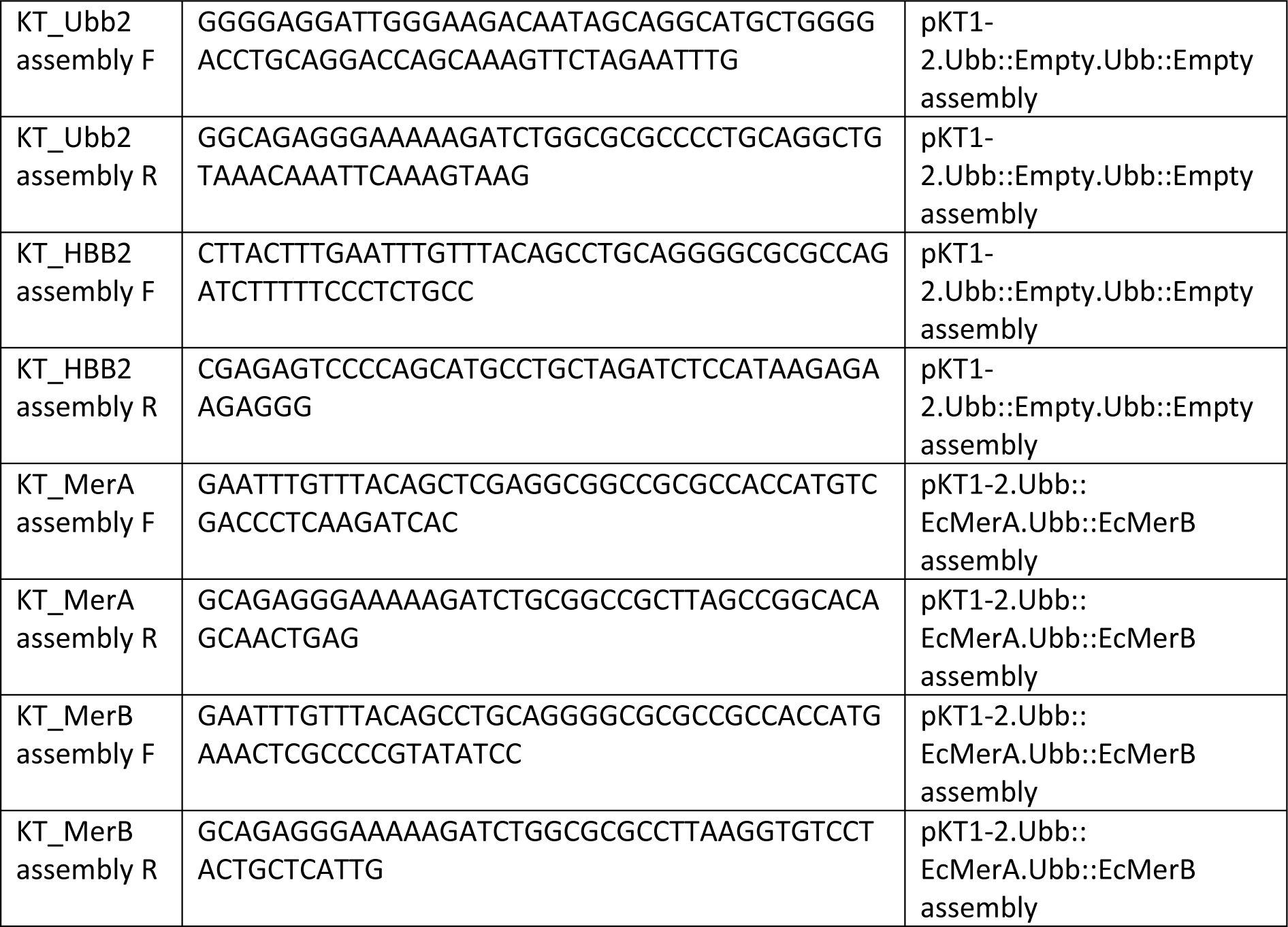
primers used in this study.

**Supplementary Table S3:**
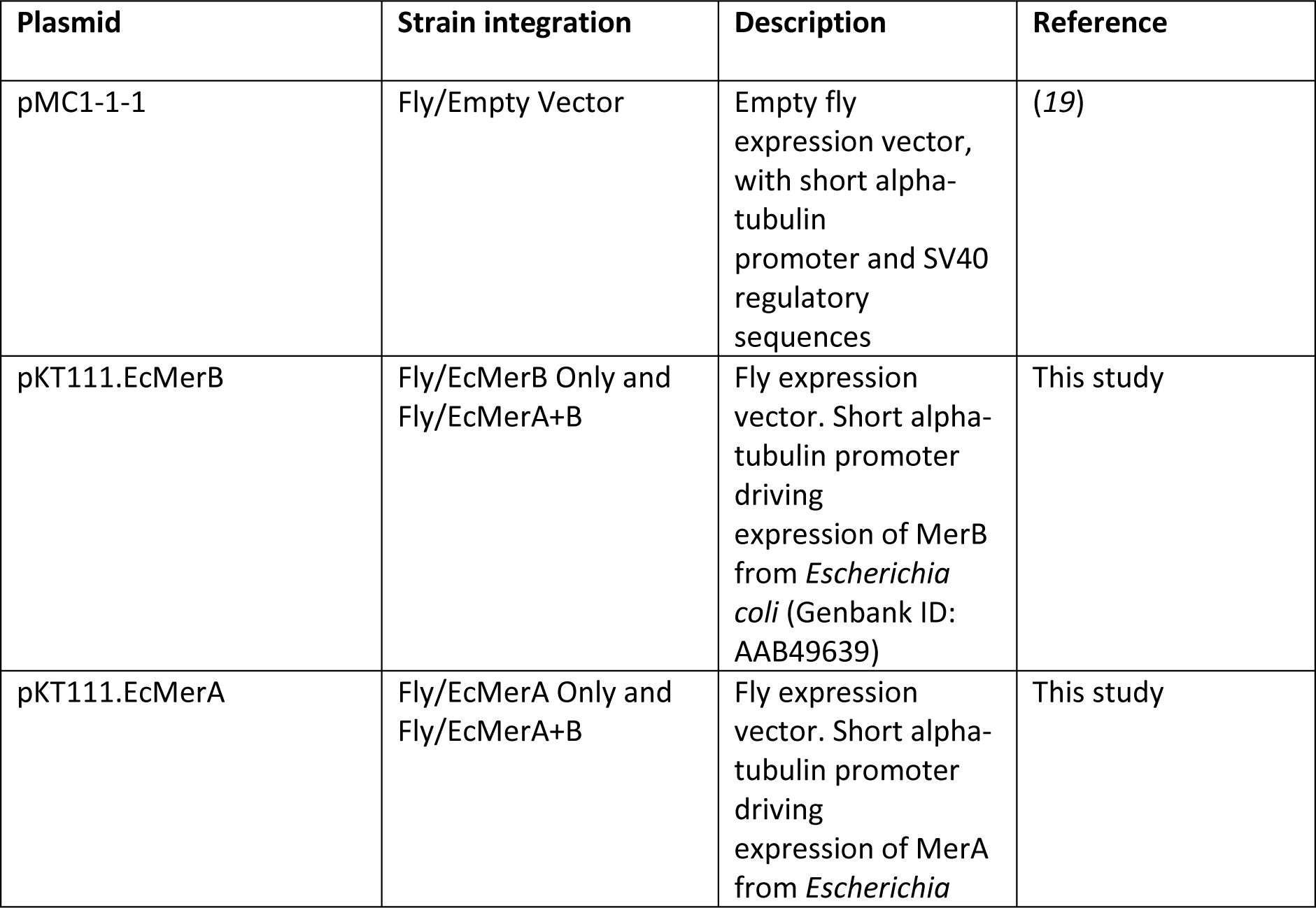
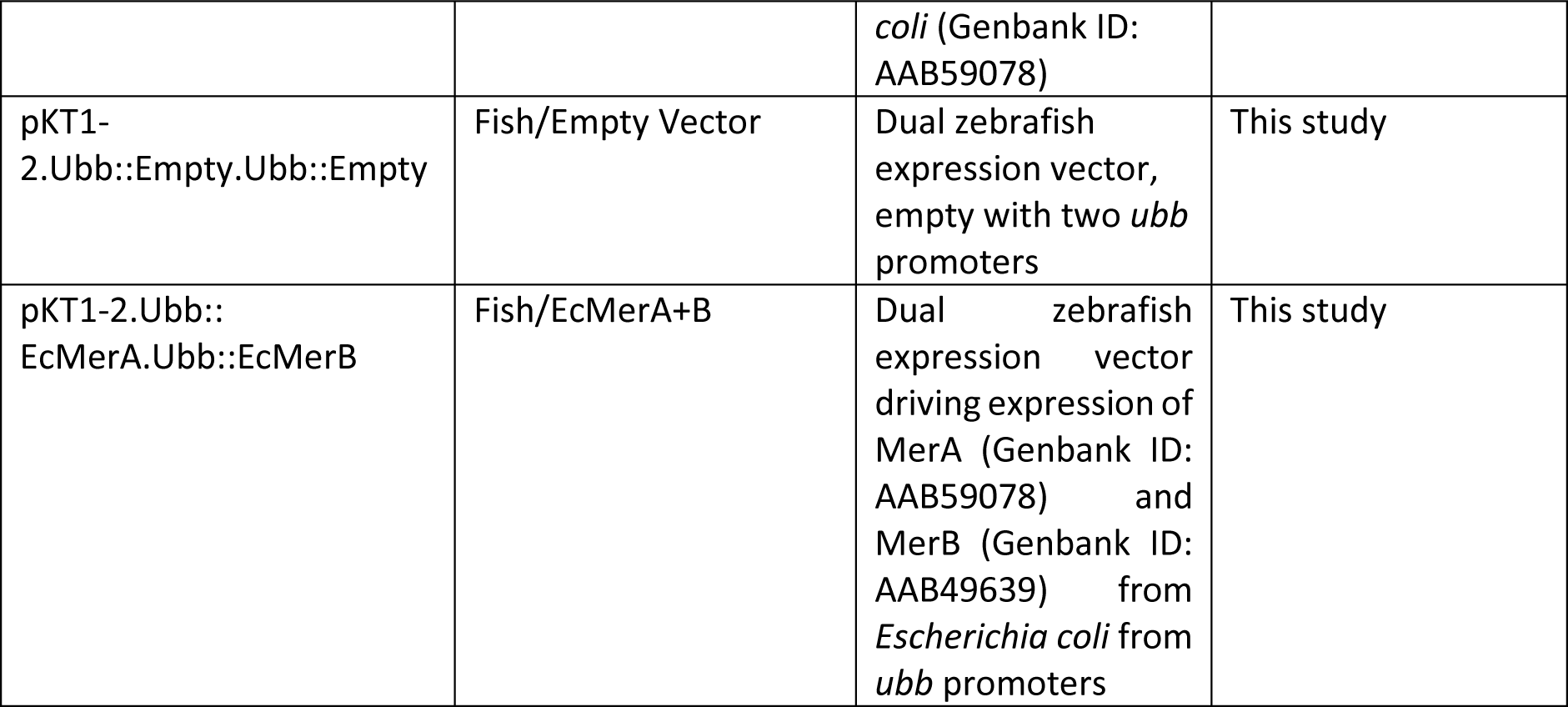
plasmids used in this study.

| Plasmid | Strain integration | Description | Reference |
| --- | --- | --- | --- |
| pMC1-1-1 | Fly/Empty Vector | Empty fly expression vector, with short alpha-tubulin promoter and SV40 regulatory sequences | (19) |
| pKT111.EcMerB | Fly/EcMerB Only and Fly/EcMerA+B | Fly expression vector. Short alpha-tubulin promoter driving expression of MerB from *Escherichia coli* (Genbank ID: AAB49639) | This study |
| pKT111.EcMerA | Fly/EcMerA Only and Fly/EcMerA+B | Fly expression vector. Short alpha-tubulin promoter driving expression of MerA from *Escherichia* *coli* (Genbank ID: AAB59078) | This study |
| pKT1-2.Ubb::Empty.Ubb::Empty | Fish/Empty Vector | Dual zebrafish expression vector, empty with two *ubb* promoters | This study |
| pKT1-2.Ubb::EcMerA.Ubb::EcMerB | Fish/EcMerA+B | Dual zebrafish expression vector driving expression of MerA (Genbank ID: AAB59078) and MerB (Genbank ID: AAB49639) from *Escherichia coli* from *ubb* promoters | This study |

**Supplementary Table S4:**
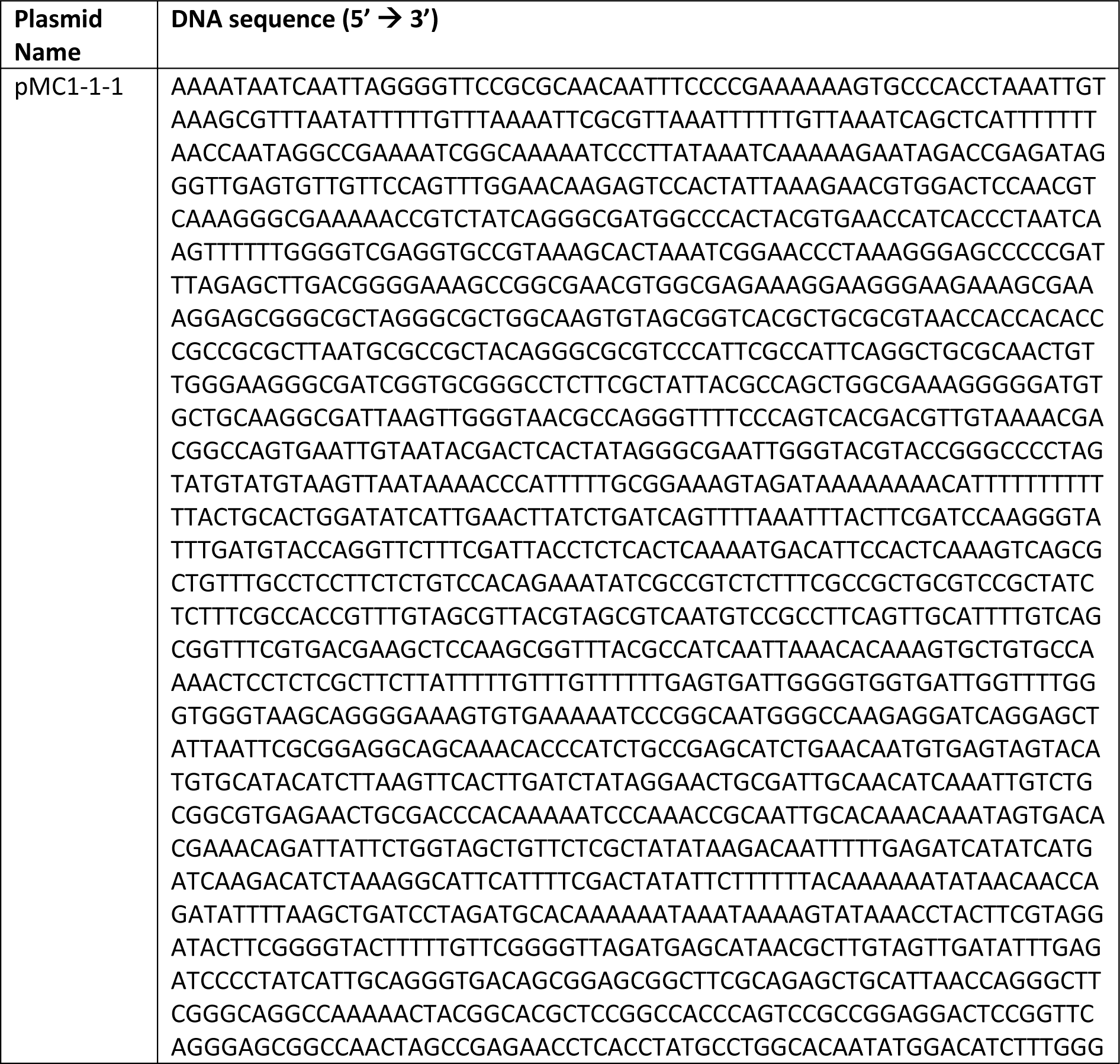
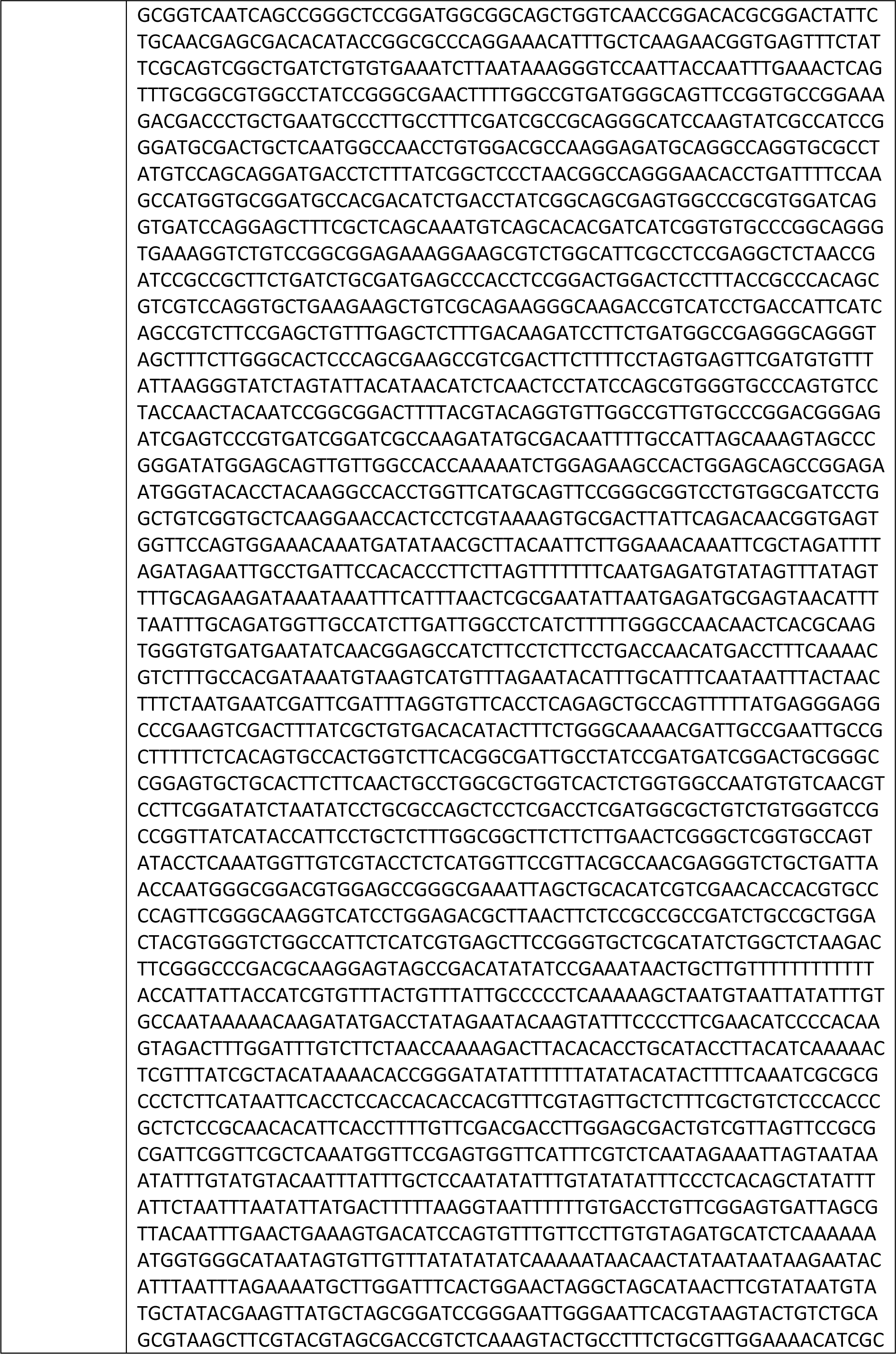
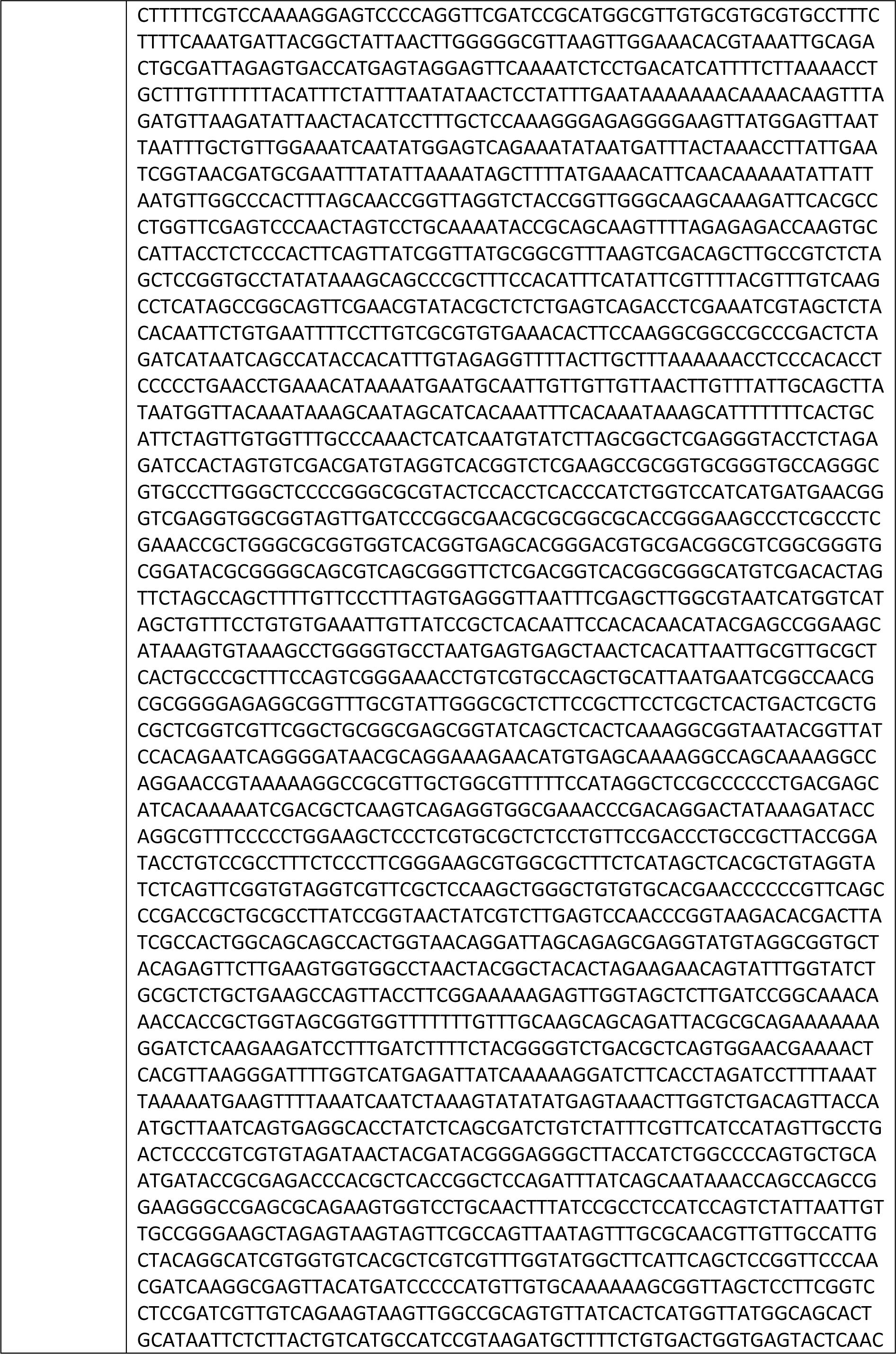
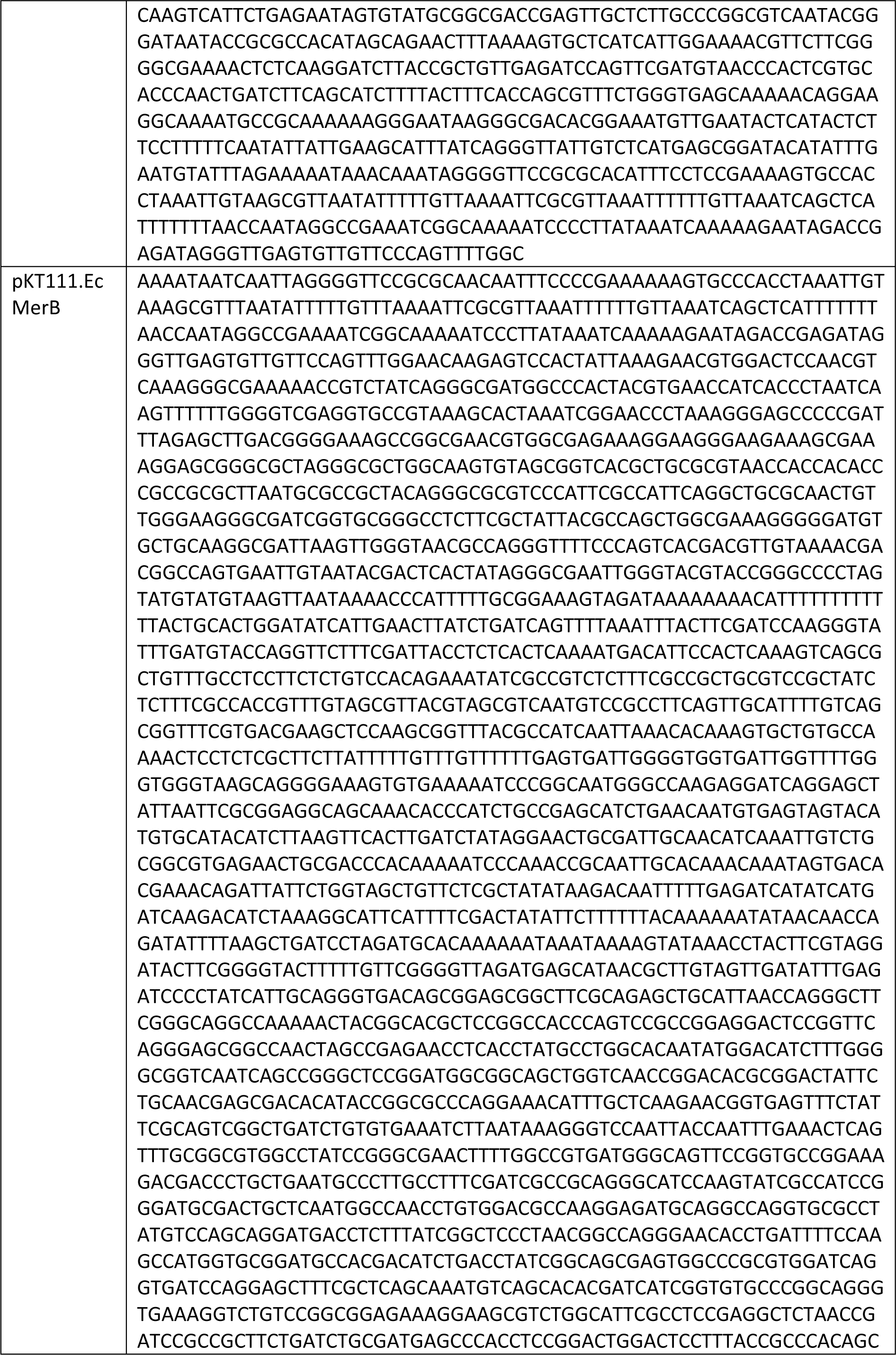
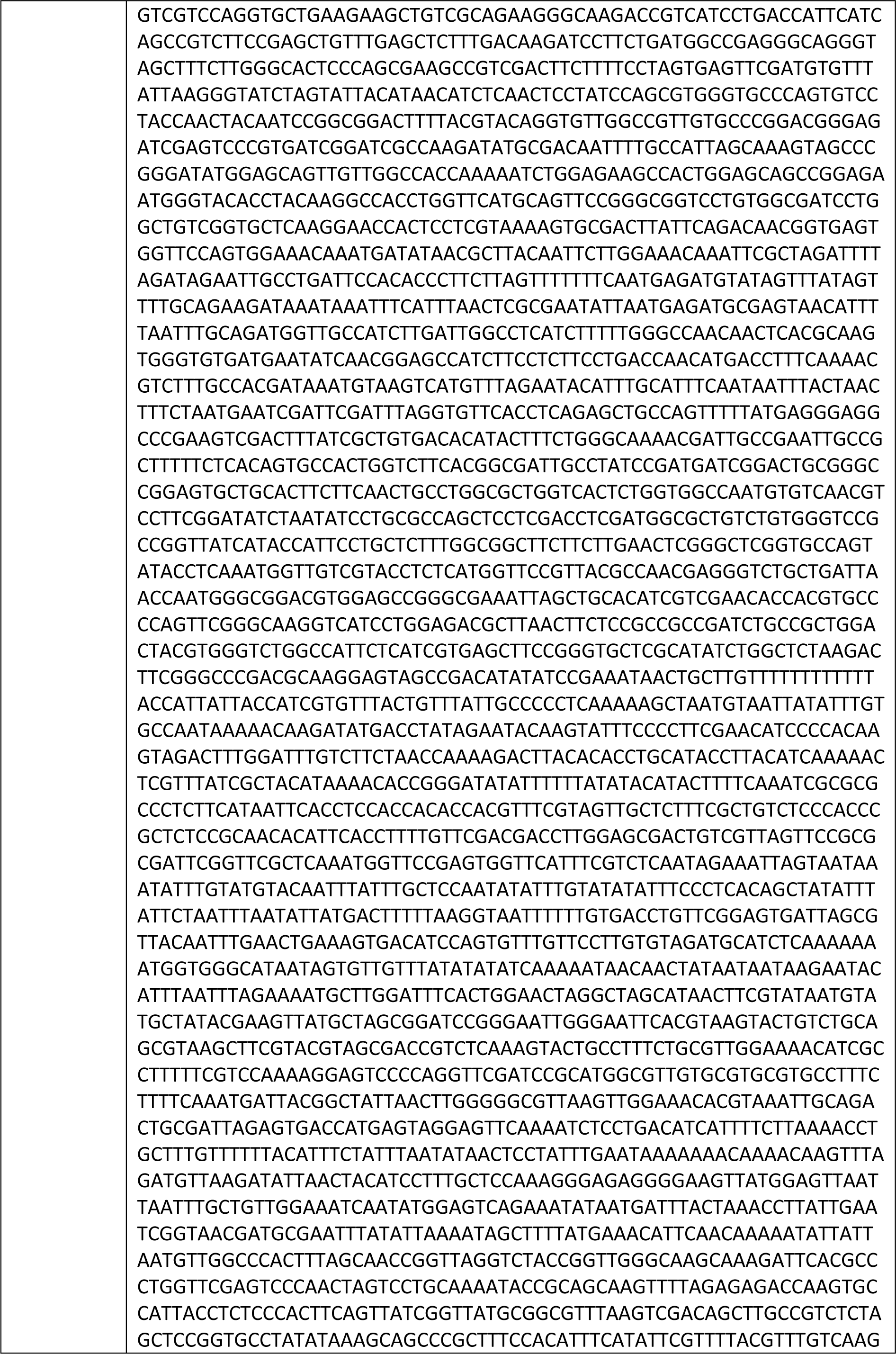
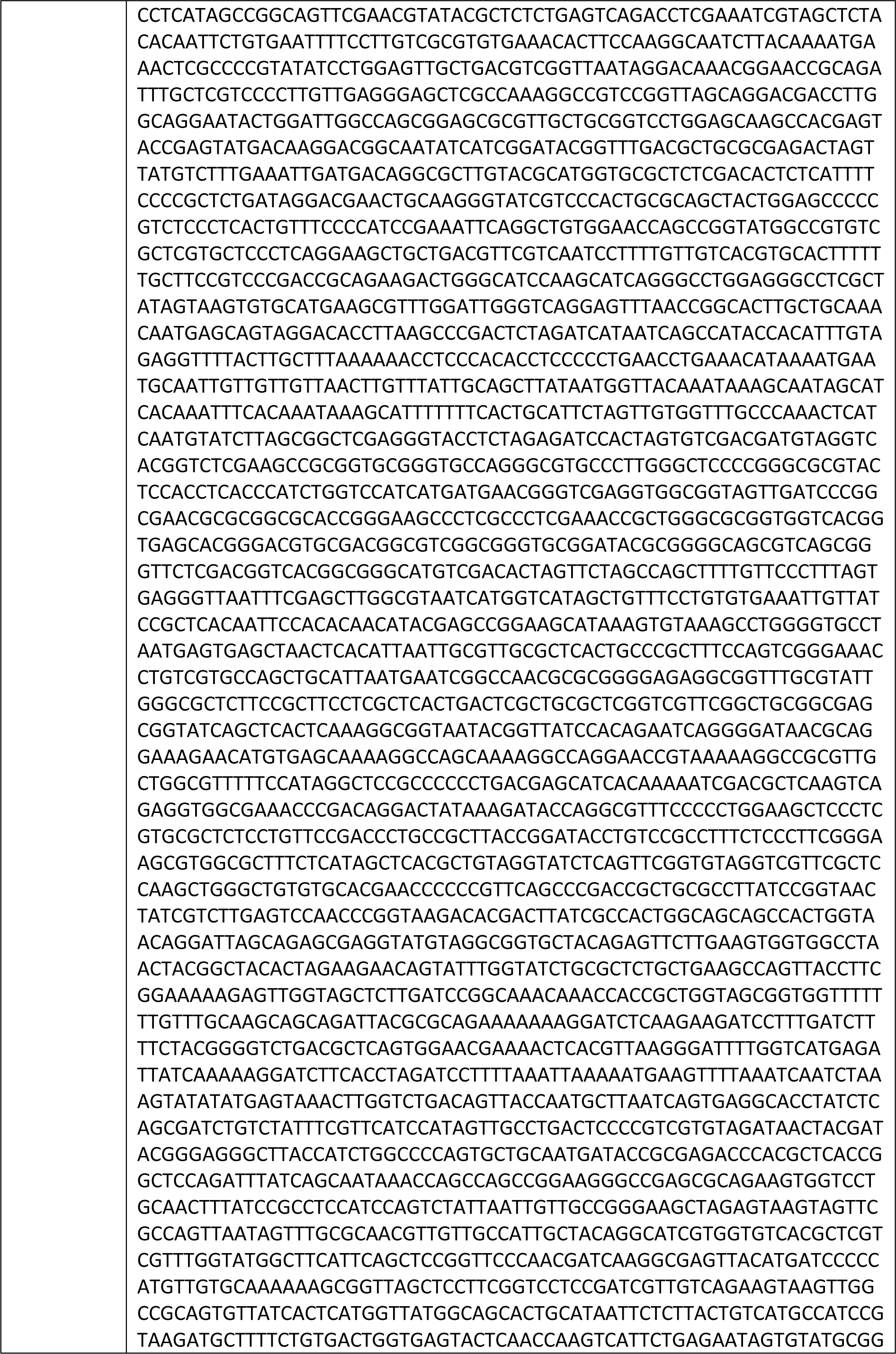
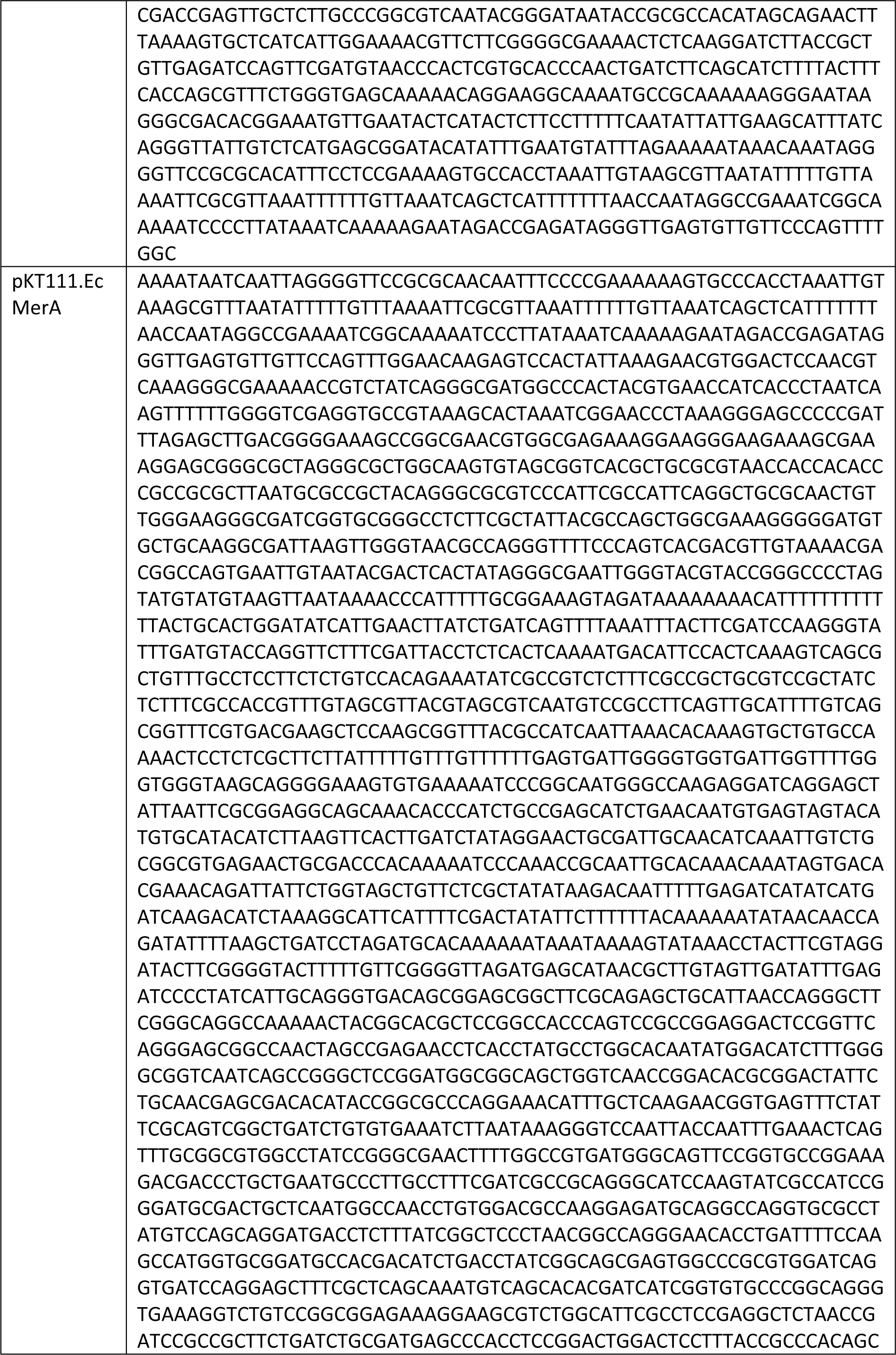
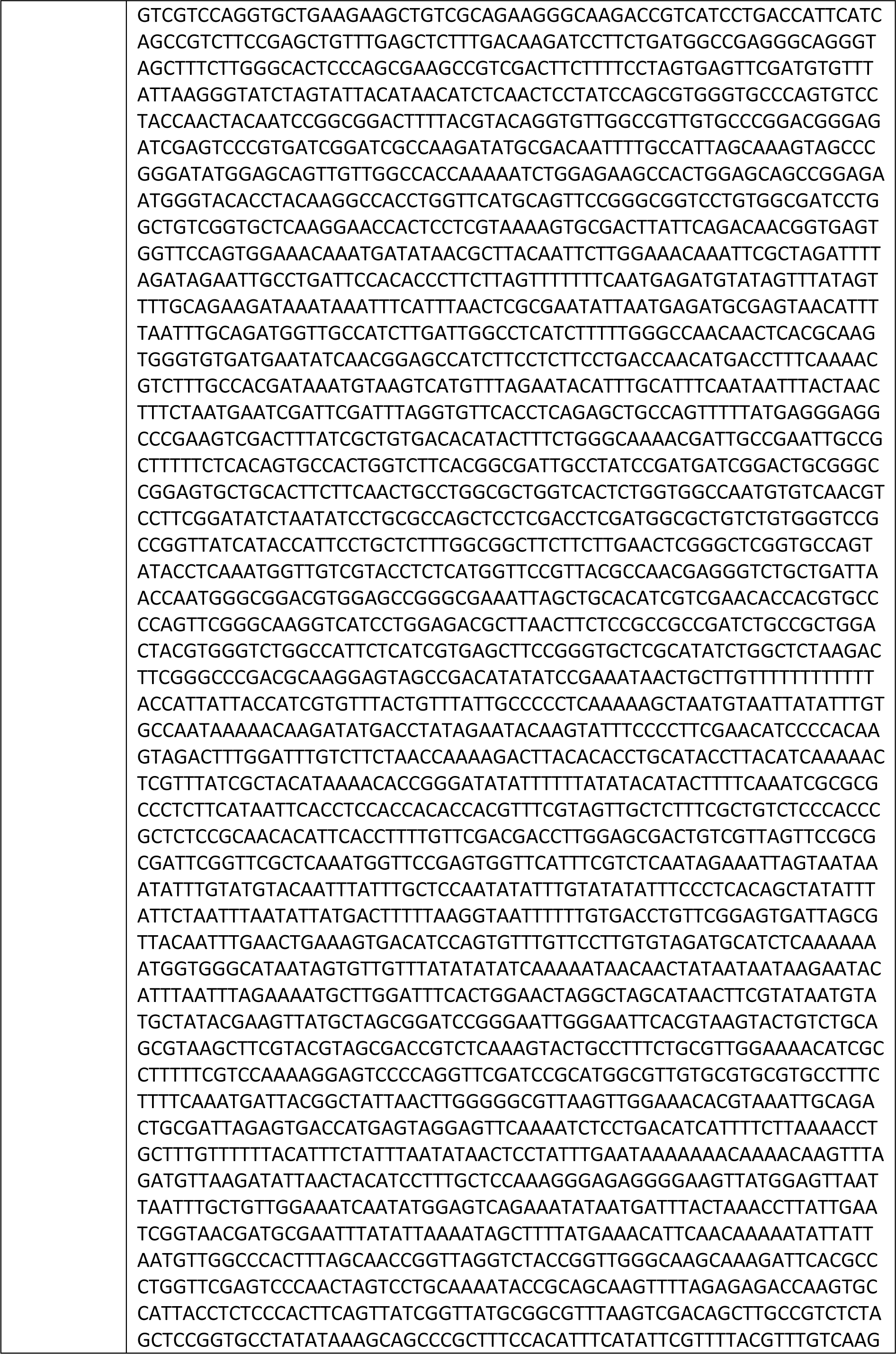
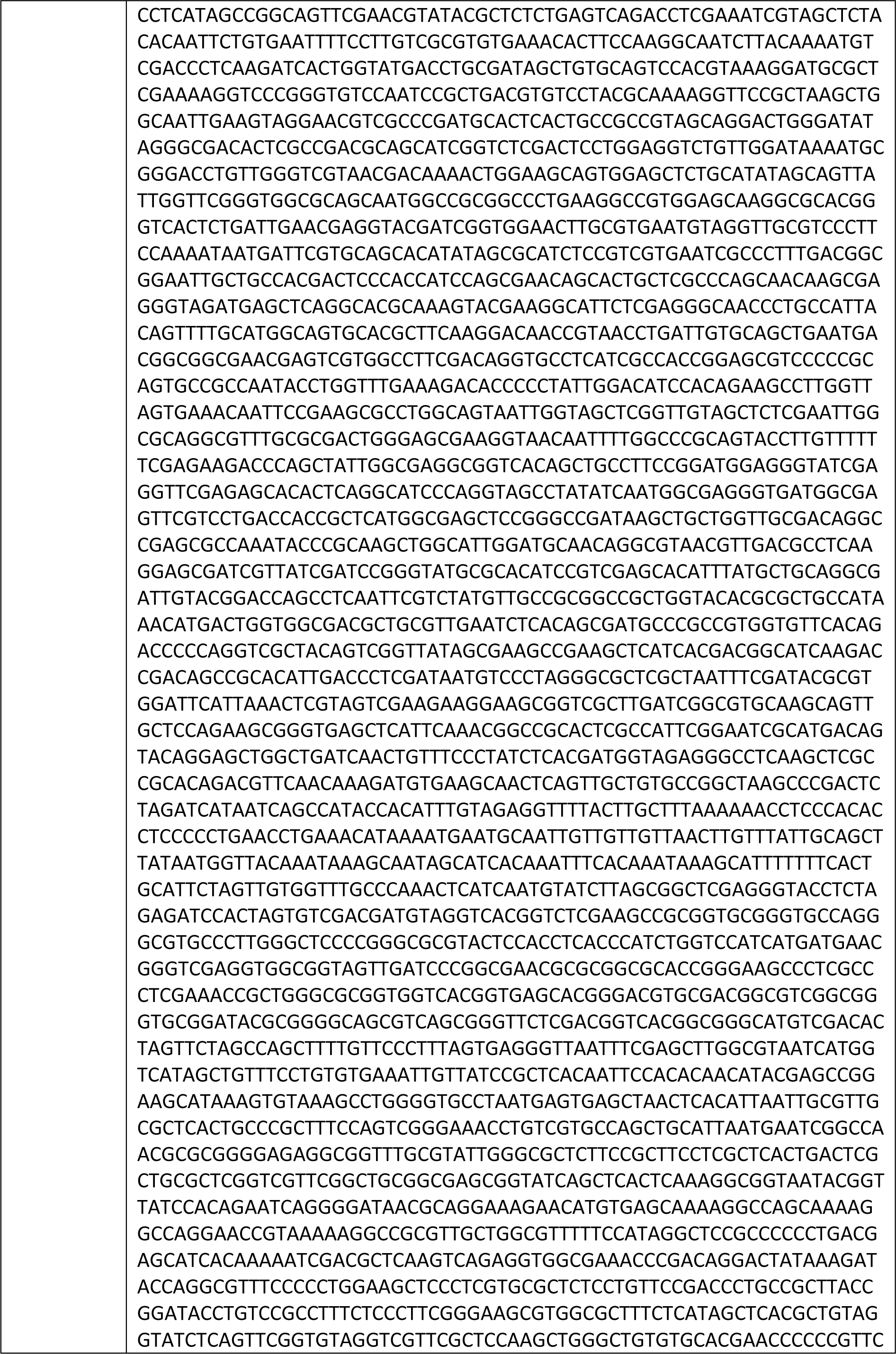
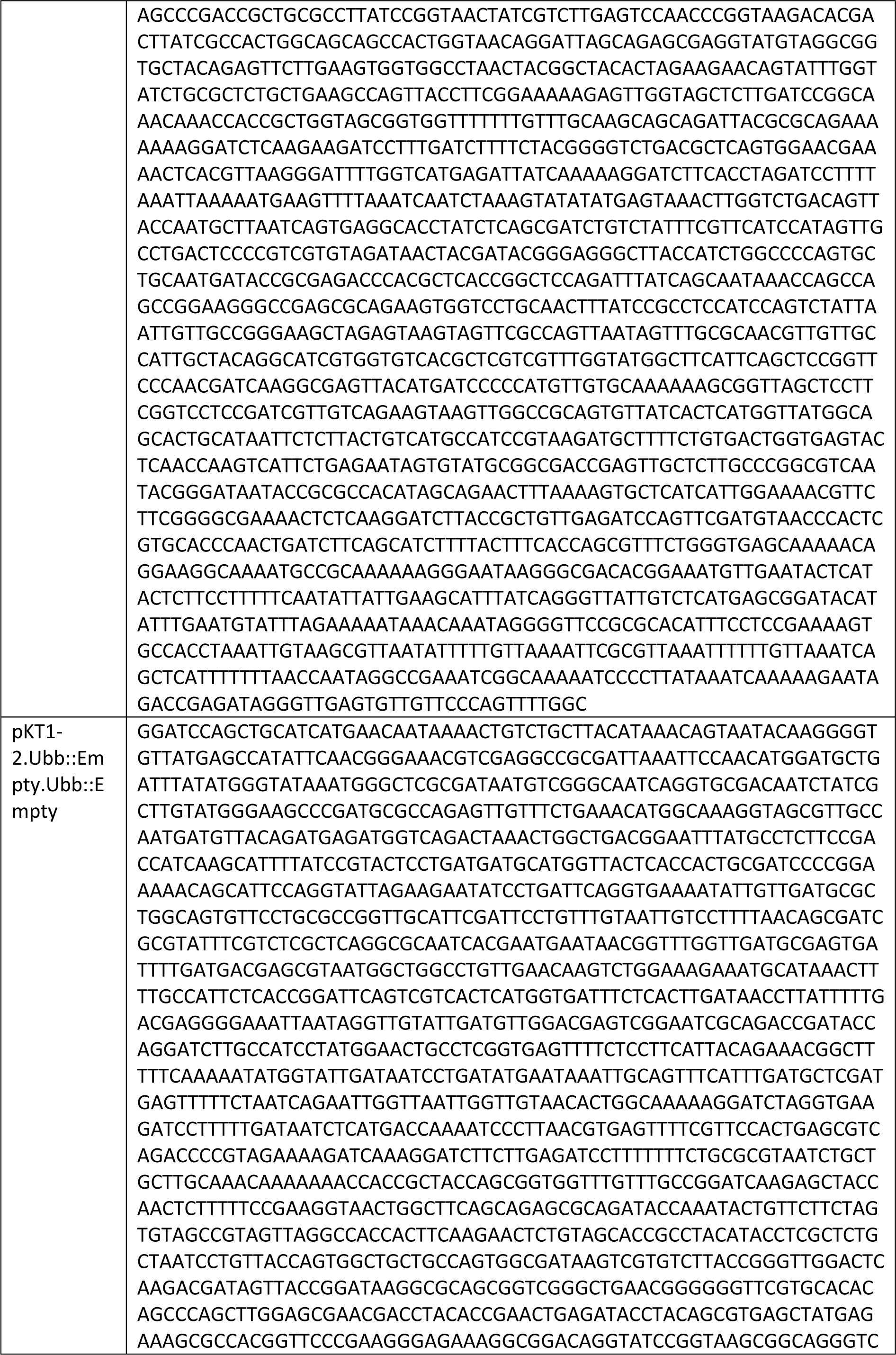
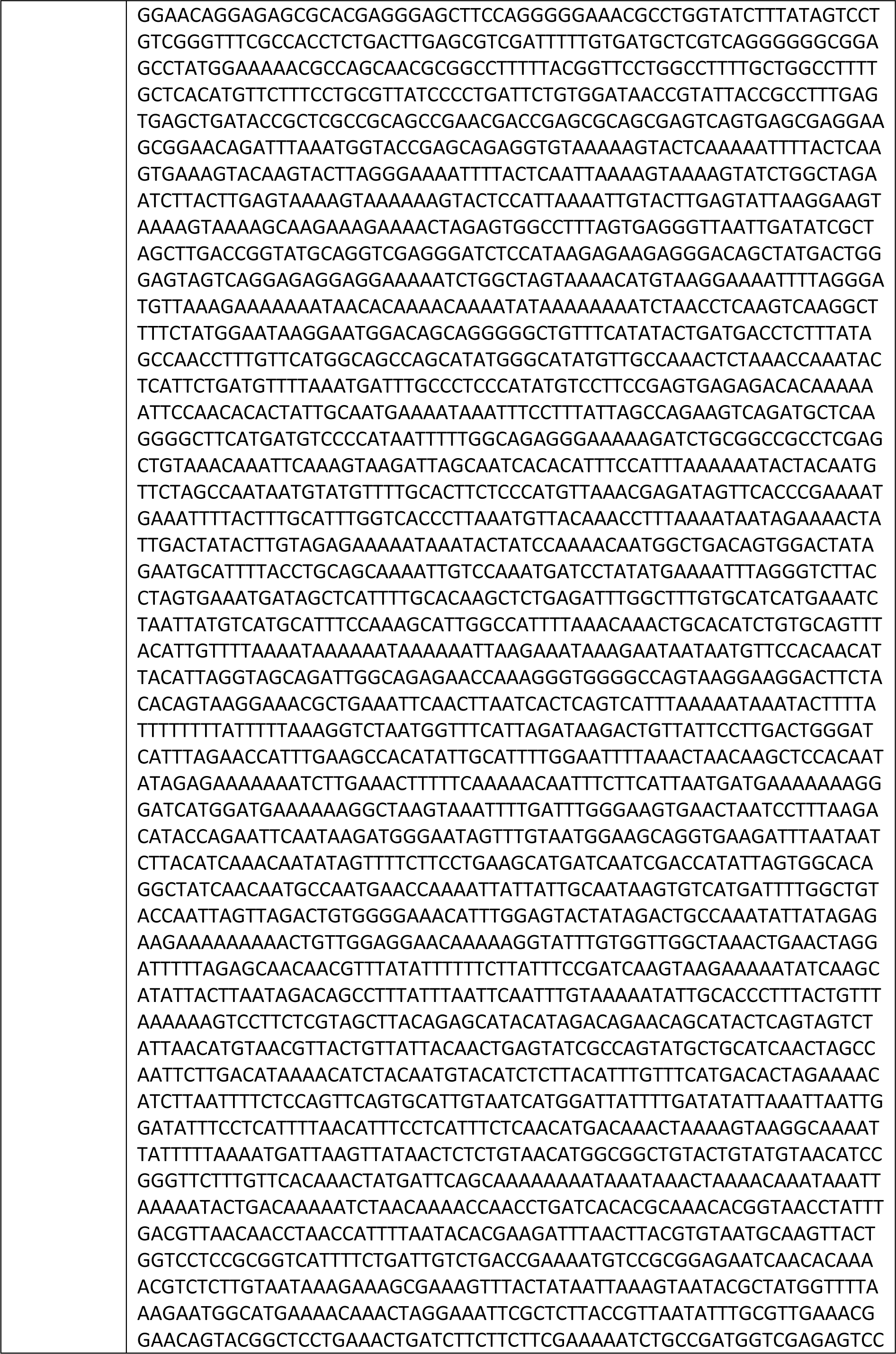
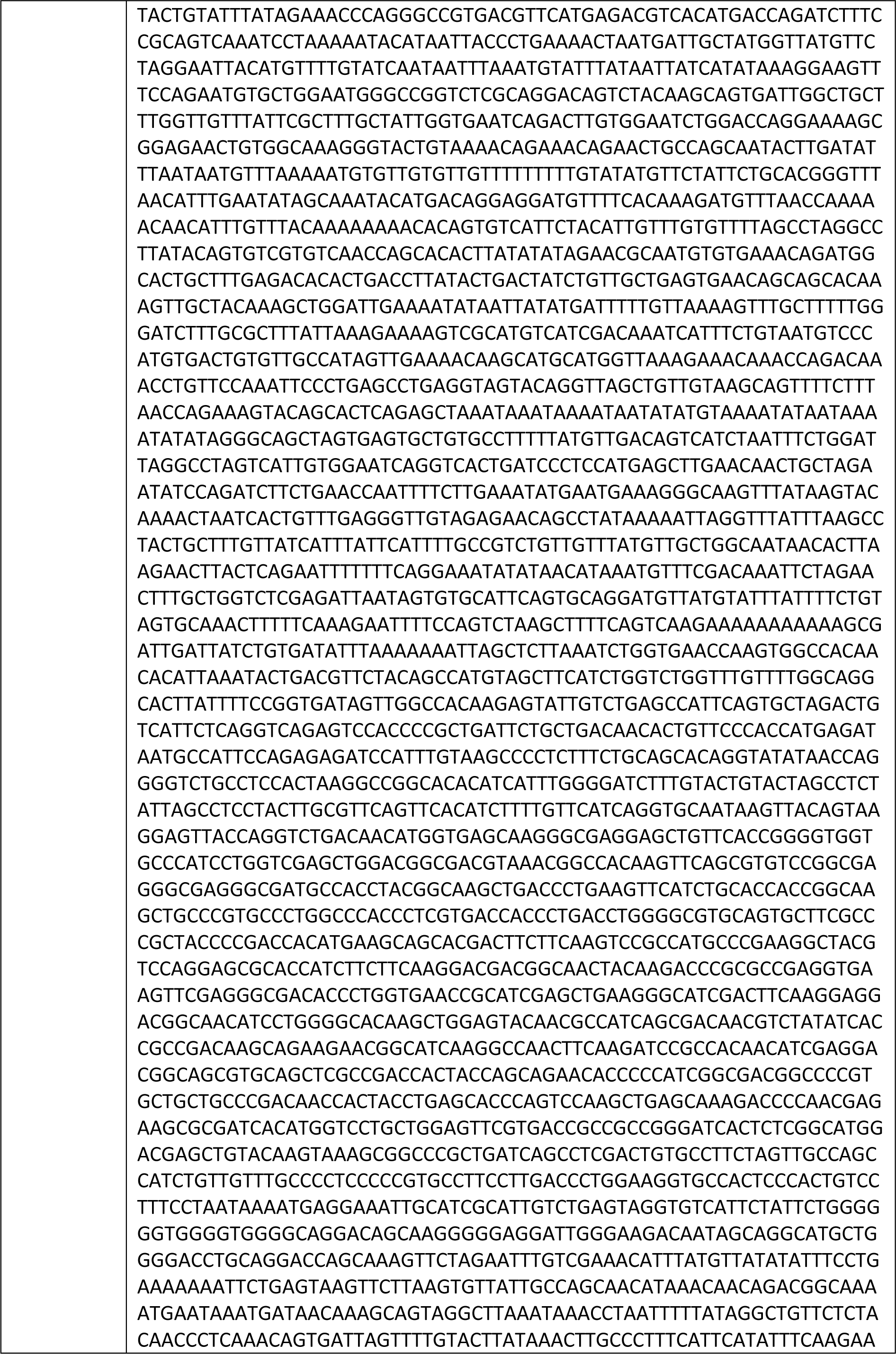
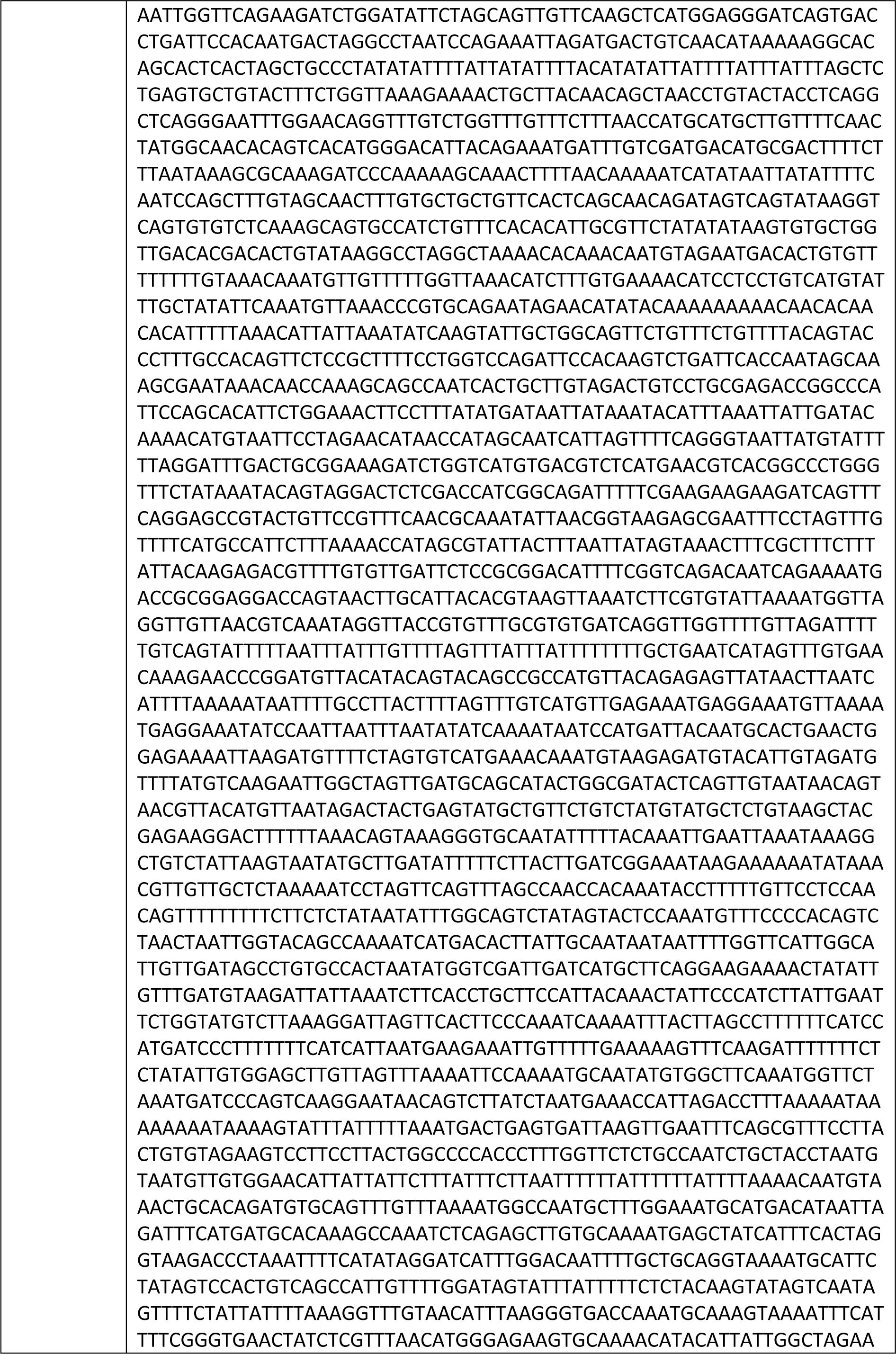
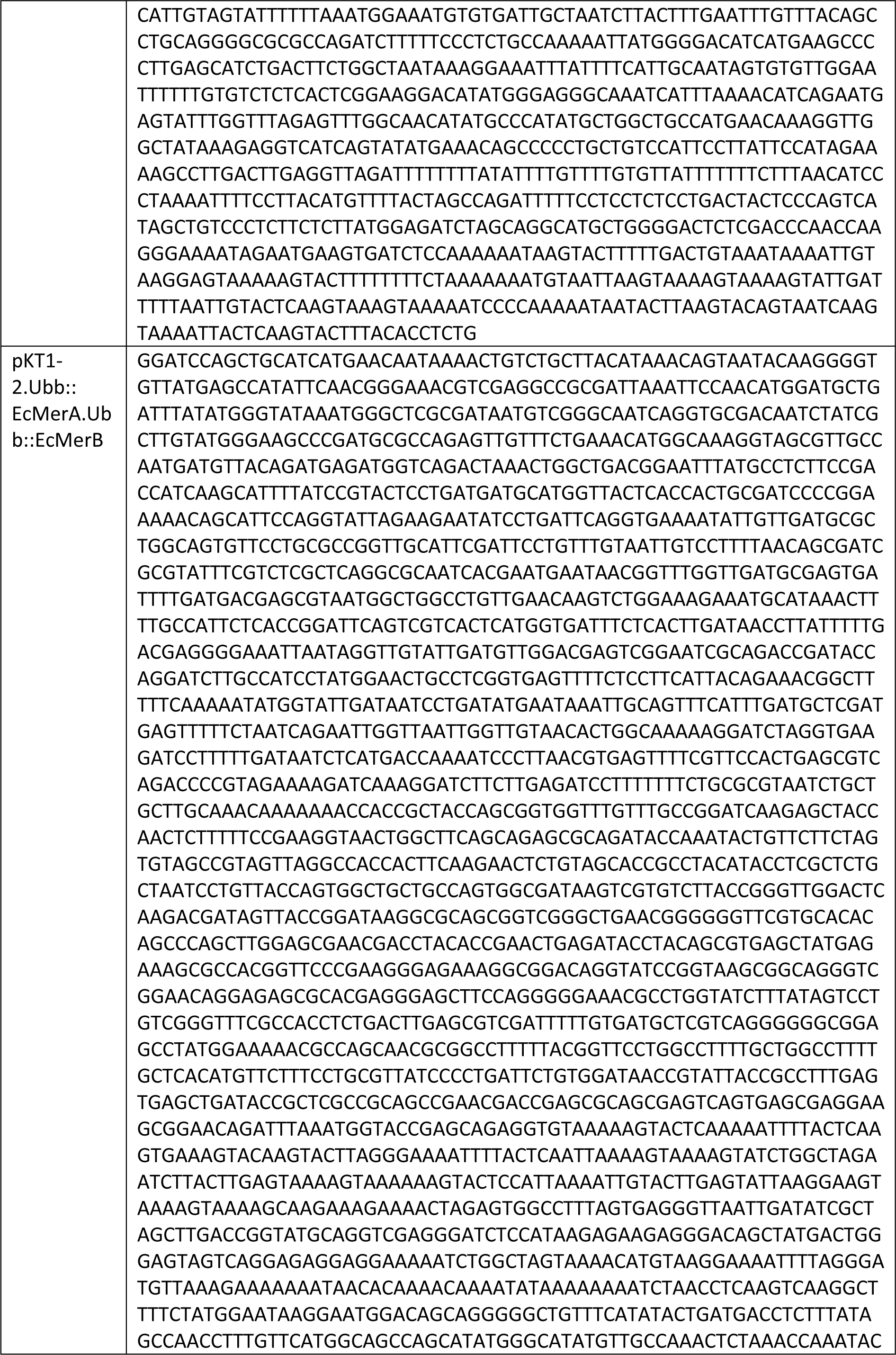
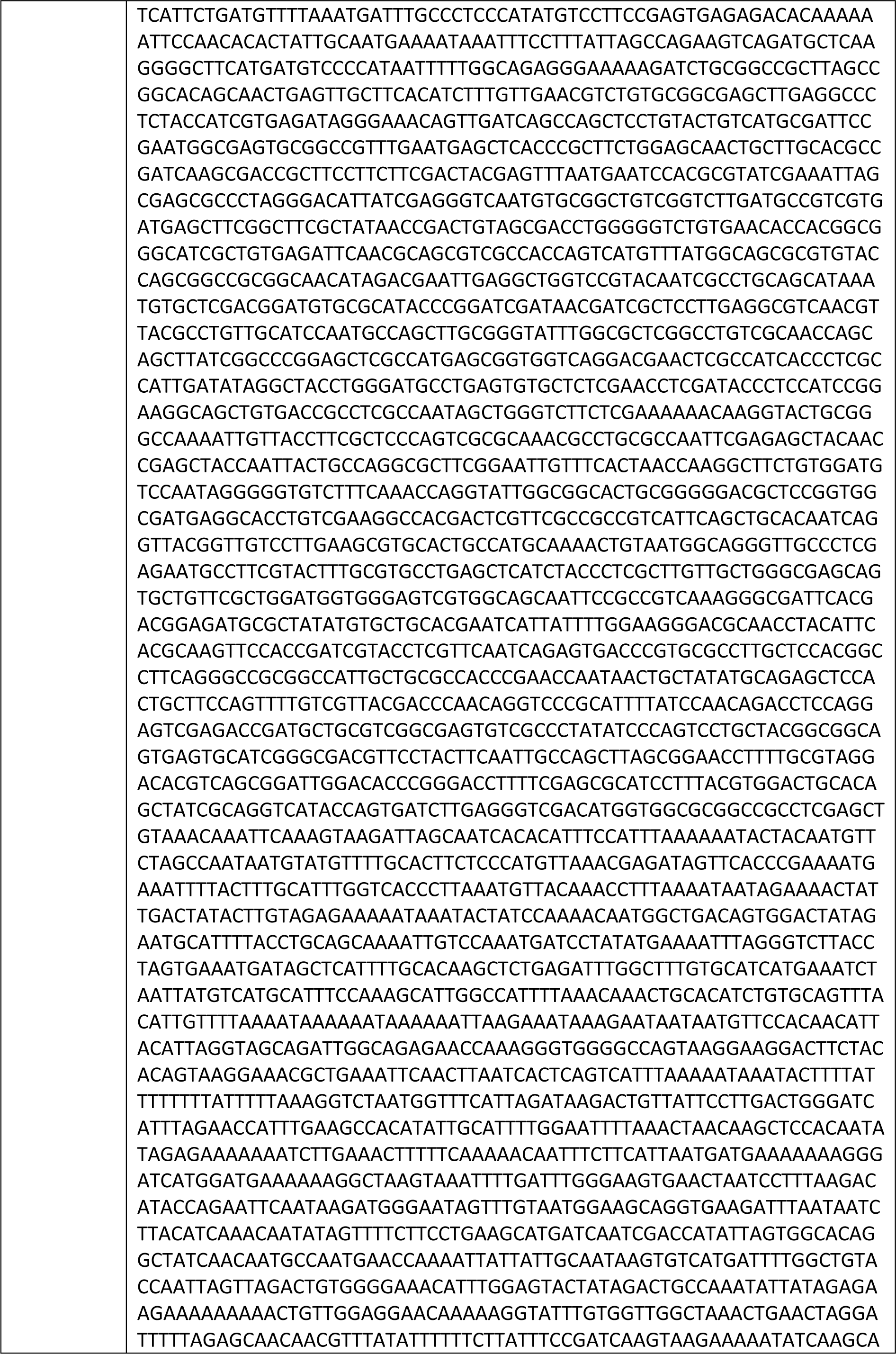
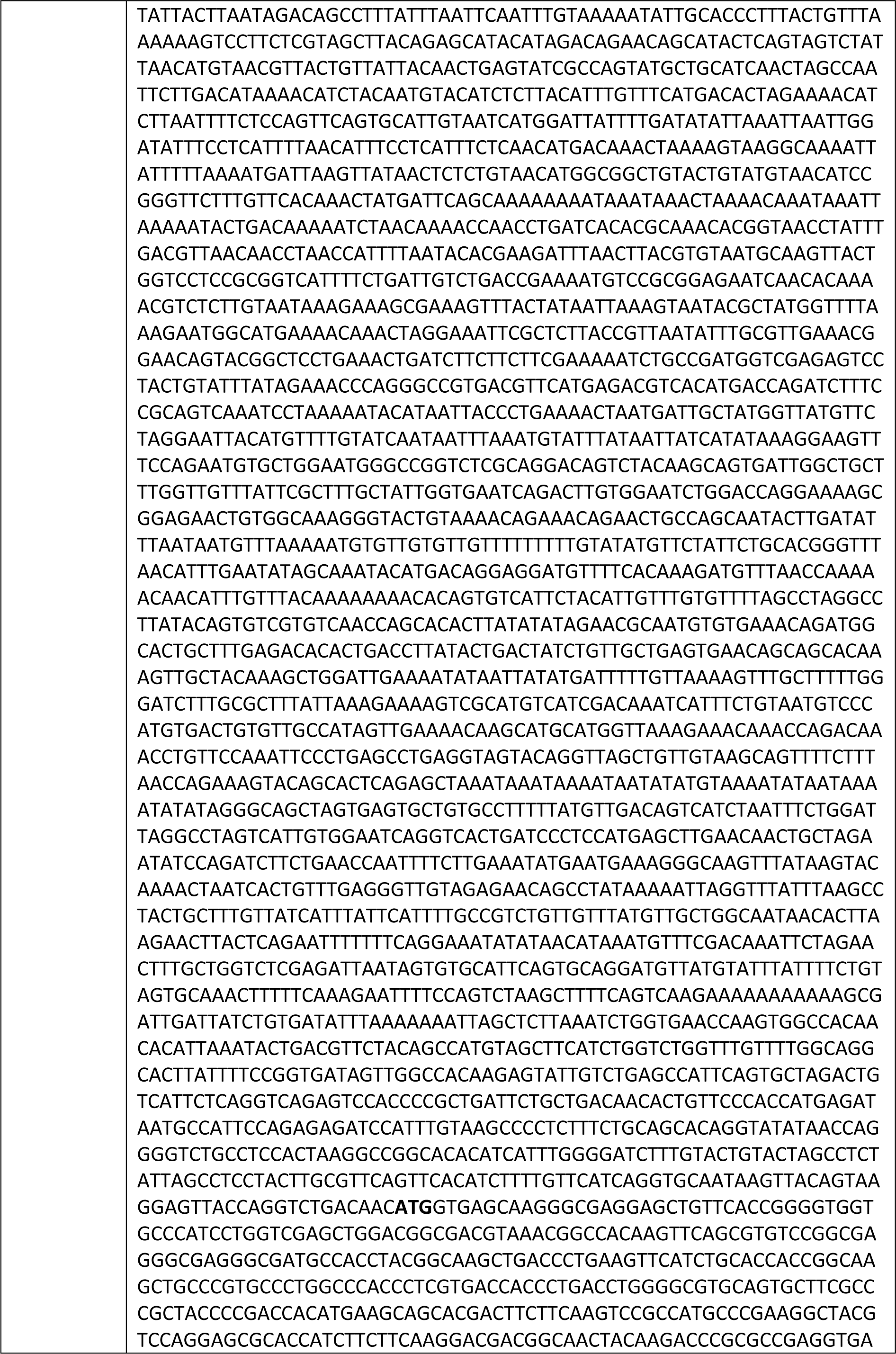
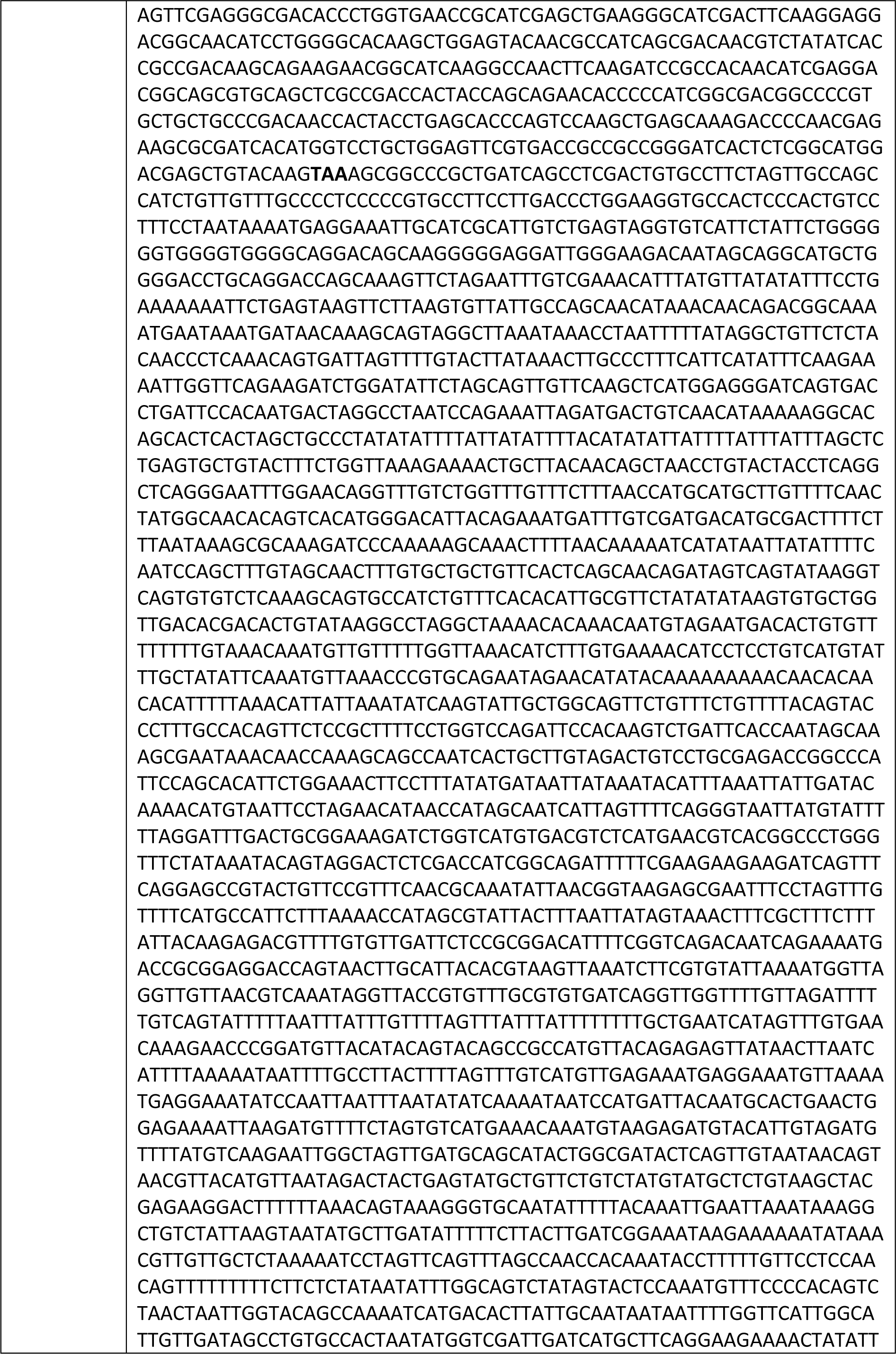
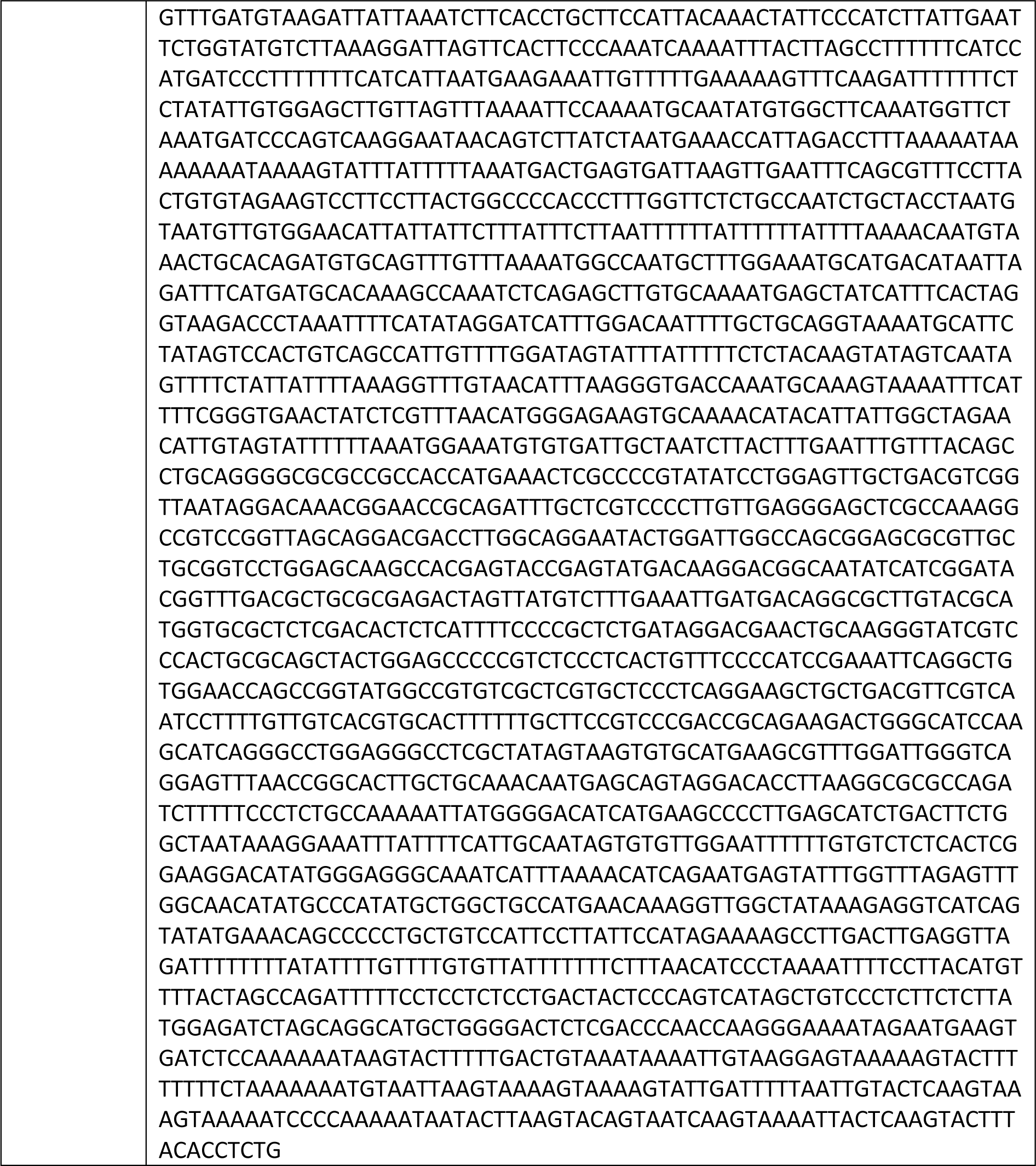
plasmid DNA sequences.

## Notes

### Competing Interest Statement

K. Tepper and M. Maselko have filed a patent application on aspects of this research.

### Summary of Updates

Revision includes citation and discussion of prior work overlooked in the original submission.

